# The Onset of Myeloid Differentiation in Hematopoietic Progenitors In Vivo is Coordinated with S-phase Progression

**DOI:** 10.64898/2026.05.10.724120

**Authors:** Jonas Metz, Alessandro Greco, Larissa Frank, Nina Claudino, Hans-Reimer Rodewald, Thomas Höfer

## Abstract

Hematopoiesis produces diverse cell types in large quantities. How hematopoietic stem cells (HSCs) initiate lineage differentiation *in vivo* is not known. Here, we jointly measure chromatin accessibility and transcriptomes in single stem and progenitor cells and uncover a tight interplay between cell-cycle progression and the onset of differentiation. The cell cycle continuously accelerates as HSCs transition to uncommitted multipotent progenitors. In these uncommitted progenitors, proliferative activation is followed by the opening of myeloid-specific chromatin loci in S phase and, subsequently, myeloid gene expression. S phase-coupled onset of myeloid differentiation becomes quantitatively enhanced in emergency myelopoiesis. At the molecular level, we find that chromatin occupancy by HOXA9 – a key transcription factor of uncommitted progenitors – predicts whether the transition through S phase will initiate myelopoiesis, or whether the uncommitted progenitor state will be retained. Taken together, our findings provide a mechanistic framework for the onset of myeloid lineage differentiation *in vivo*.

## INTRODUCTION

Hematopoiesis is a paradigm for the continuous generation of diverse cell types from a tissue stem cell. How a hematopoietic stem cell (HSC) gives rise to diverse cell fates has long been a key question. Early pioneering work, based on transplantation of sorted progenitors in mice, identified oligolineage progenitor populations that had lost some lineage fates but retained several others^1,2^. Subsequently, single-cell transcriptomics introduced the notion that multipotent progenitors (MPPs) and common myeloid progenitors (CMPs) constitute a mixture of progenitor states at the single-cell level^3–8^. Moreover, barcoding and fate-mapping of phenotypically defined HSCs have shown that HSC clones themselves may give rise to all lineages or be restricted to erythro-myeloid fates^9–11^, including clones with strong production of platelets^12,13^. Collectively, these studies define different routes from distinct stem cell populations to mature blood and immune cell types. However, they leave open the fundamental question of how cells exit the stem cell state and initiate lineage decisions in native hematopoiesis.

Transcription factors specific to hematopoietic lineages, as well as cytokines that can instruct lineage choice in cell culture^14^ have been identified. Moreover, asymmetric cell division has been implicated in HSC differentiation in culture^15^. *In vivo*, the production of mature cells is indeed regulated by cytokines. The regulation of erythropoiesis and thrombopoiesis by, respectively, erythropoietin and thrombopoietin are prominent examples^16,17^. These mechanisms act after terminal lineage choices to erythrocytes or platelets have already been made and regulate the rate of mature cell production. However, progenitors with restricted fates emerge from stem cells upstream of these homeostatic regulatory mechanisms^7,10^. A robust mechanism for initiating stem cell differentiation in native hematopoiesis is missing.

A characteristic feature of hematopoiesis is its enormous productivity. In the adult mouse, approximately 1% of all HSCs, of the order of 10^2^ cells, differentiate per day as the ultimate source of the production of several times 10^8^ mature blood cells^18^. Hence, about several million hematopoietic progenitor cell divisions take place every day^19^, with progenitors dividing at least an order of magnitude more frequently than HSCs^20^. Indeed, proliferative multipotent progenitors are the central source of the day-to-day replenishment of blood cells^17^. For committed erythroid progenitors, it has been shown that the cell cycle itself, specifically an accelerated S phase, is required for differentiation^21,22^. These and similar observations^23–25^ pose the question whether upstream of these specific lineage pathways, in uncommitted progenitors, cell cycle acceleration and differentiation are linked.

Here we address this question by assaying simultaneously chromatin accessibility and transcriptomes in single hematopoietic stem and progenitor cells (HSPCs). This single-cell multiome approach has recently emerged as a powerful method to identify regulatory changes at the chromatin level in cell differentiation^26,27^. Here, we develop a computational framework to interrogate the interplay between cell differentiation and cell-cycle progression. Thus, we discover a robust cell-cycle signature in chromatin accessibility – a feature that has previously been thought to be inert to the cell cycle. This signature allows us to probe systematically lineage gene opening in relation to cell-cycle phase. We find that cell-cycle acceleration precedes the opening of myeloid and erythroid gene-regulatory regions, subsequently giving rise to differentiating progenitors. In uncommitted progenitors, myeloid-specific chromatin opens in S phase, suggesting that proliferative activation initiates lineage differentiation. Indeed, in emergency myelopoiesis triggered by systemic infection, progenitor proliferation is accelerated and the coupling between S phase progression and myeloid differentiation becomes strongly enhanced. At the molecular level, we find that high chromatin occupancy by HOXA9 in uncommitted HSPCs predicts that after S phase transition the stem-cell-like state will be retained, whereas low HOXA9 occupancy predicts the initiation of myelopoiesis during DNA replication.

## RESULTS

### Proliferative activation of progenitors precedes myeloid differentiation

To produce the multitude of hematopoietic cell types at the required numbers, HSC activation results in both differentiation and enhanced cell proliferation. To probe these processes during steady-state hematopoiesis, we profiled transcriptomes and open chromatin simultaneously in single HSPCs freshly isolated ex vivo in three independent experiments (Figure 1A). Changes in chromatin accessibility report on regulatory events preceding lineage gene expression, while the transcriptome reports on lineage gene transcription and cell-cycle state. To probe these molecular features in individual HSPCs, we sorted, from the bone marrow of wildtype mice, primitive (Lin^−^Sca1^+^Kit^+^) populations, specifically HSCs (CD48^−^ CD150^+^), ST-HSCs (CD48^−^ CD150^−^) and MPPs (CD48^+^ CD150^−^), as well as lineage-committed progenitors, Lin^−^Kit^+^ (LK) cells. Single nuclei were extracted and both RNAseq and ATACseq were performed. After quality control, we obtained a total of 17,604 HSPCs for joint analysis of chromatin accessibility and transcriptomes, without the need for computational alignment (2003 HSCs, 1988 ST-HSCs, 5425 MPPs and 8188 LK cells combined from the three mice).

**Figure 1.**
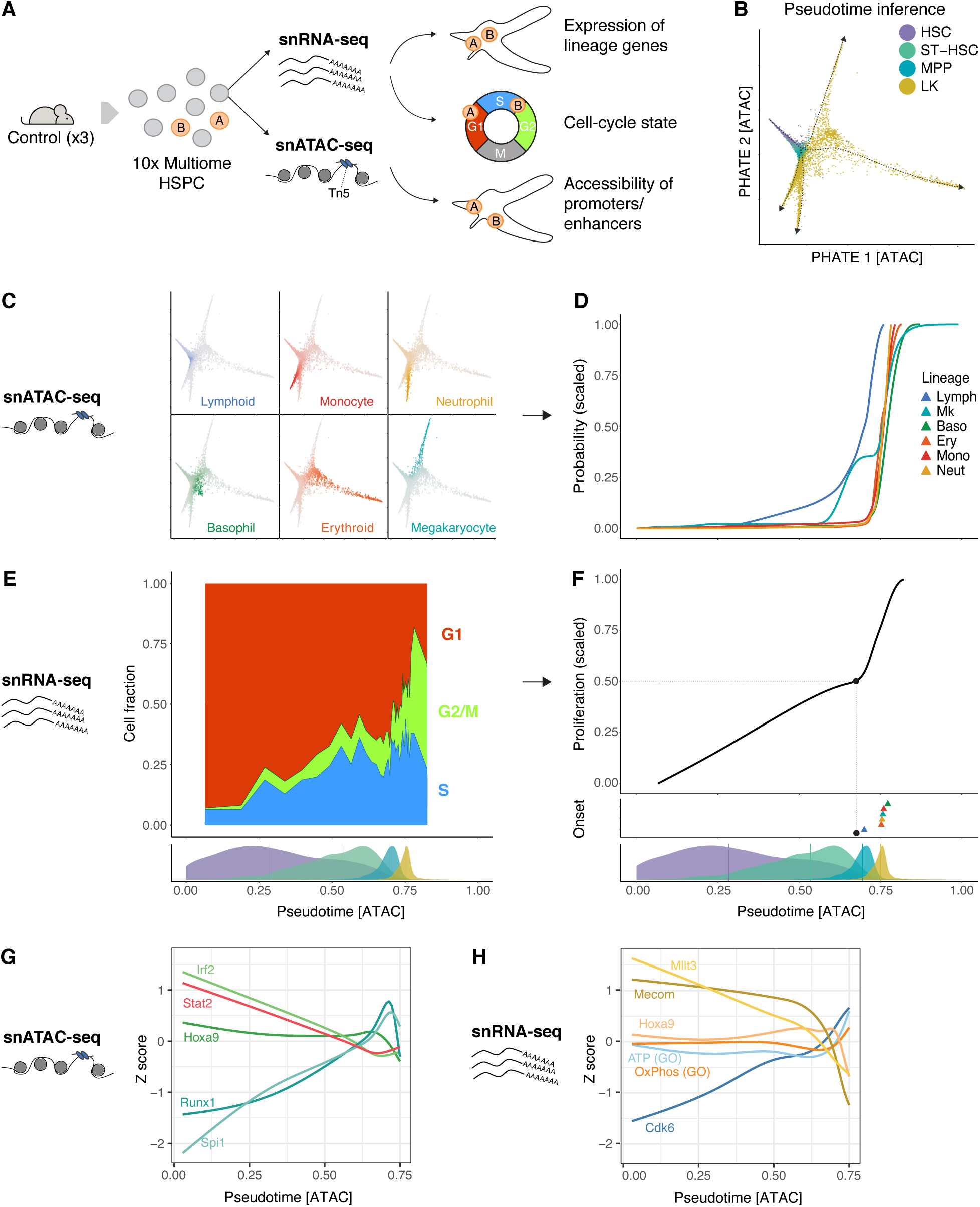
Proliferative activation precedes hematopoietic lineage differentiation. (A) Schematic of the experimental approach. Lin^−^Sca1^+/-^Kit^+^ cells were isolated from the bone marrow of untreated wildtype C57BL/6J mice (3 biological replicates). These cells were subjected to simultaneous single-nucleus RNA sequencing (snRNA-seq) and single-nucleus ATAC sequencing (snATACseq). Cell-cycle phase was inferred using transcripts, while the differentiation state was inferred using both transcriptome and open chromatin. (B) PHATE visualization based on chromatin accessibility at hematopoietic enhancers. Cells are colored based on the FACS gate. (C) Single cells colored based on fate probability inferred by CellRank on major lineages detected in the dataset. (D) Lineage probabilities from (C) as a function of pseudotime. (E) Fraction of cells in S, G2/M, and G1 phases of the cell cycle as a function of pseudotime. The small panel at the bottom indicates the distribution of sorted cells, HSCs, ST-HSCs, MPPs and LK cells (colorcode as in B). (F) Moving average of the fraction of cells that are not in G1 phase. The small panel at the bottom indicates the distribution of sorted cells, HSCs, ST-HSCs, MPPs and LK cells (colorcode as in B). (G) Accessibility dynamics at transcription factor binding motifs during proliferative activation of hematopoietic stem cells (ChromVar z-scores). (H) Dynamics of mRNA counts for single genes and gene modules (counts are quantified as z-scores). (B)-(H) Data from Experiment 1, other replicates are shown in Figure S1.

To identify chromatin changes underlying lineage specification, we embedded cells by their chromatin accessibility at hematopoietic enhancers ^28^, using algorithms designed to preserve the nonlinear geometry of differentiation landscape (using the diffusion map^29^ for analyses and PHATE^30^ for low-dimensional visualization). This revealed a regulatory landscape of hematopoiesis, where an uncommitted population at the tip, containing HSCs, ST-HSCs and part of the MPPs, splits into several branches, comprising the remaining MPPs and LK cells (Figure 1B). On the basis of this enhancer-based differentiation landscape, we computed diffusion pseudotime values for all cells, which was consistent with a differentiation progression from the stem-cell state towards multiple lineage branches (Figure 1B, indicated by arrows). Assigning enhancer-based lineage probabilities to individual cells with CellRank^31^ identified major differentiation trajectories to megakaryocytes, erythrocytes, lymphocytes, and myeloid lineages (neutrophils, monocytes, and basophils) (Figure 1C). The analysis of lineage gene opening versus transcript levels showed that chromatin changes foreshadowed transcription (Figure S1). Thus, we obtained a chromatin-based differentiation landscape of hematopoiesis where regulatory events of lineage specification in the chromatin precede lineage gene expression.

We exploited the joint information on cell cycle genes and opening of lineage enhancers to determine the relation of proliferation onset and lineage specification. For erythroid and myeloid (basophil, monocyte, neutrophil) differentiation, lineage specification occurred abruptly at similar pseudotime in these lineages (Figure 1D; corresponding to late MPPs, bottom pseudotime panel Figure 1F), whereas lymphoid and megakaryocyte chromatin features began to appear earlier (Figure 1D). Cell-cycle phases, based on transcripts, showed an incremental increase in the cycling fraction in stem cells, followed by an abrupt rise in progenitors (MPPs and LK cells) (Figure 1E), consistent with pulse-chase experiments^32,33^. The rise in the fraction of cycling cells preceded differentiation for neutrophil-monocyte, basophil, and erythrocyte lineages, and to a lesser extent also megakaryocytes (Figure 1F, Experiment 1; Figure S2, the other two biological replicates). Thus, proliferative activation precedes myeloid and megakaryocyte-erythroid differentiation.

Next, we probed molecular changes prior to lineage commitment. Transcription factor (TF) motifs related to hematopoietic progenitor specification (RUNX1 and PU.1, encoded by *Spi1*) progressively opened as stem cells differentiated to MPPs (Figure 1G). By contrast, binding motifs for HOXA9, which is selectively expressed in uncommitted hematopoietic cells, remained at a plateau in stem cells and only dropped sharply at the transition from MPPs to LKs. Motifs for interferon response TFs (STAT2 and IRF2) progressively closed, consistent with intrinsic expression of interferon-stimulated genes selectively in stem cells^34^ (Figure 1G).

Identifying transcripts that changed as HSCs differentiated to progenitors, we found a progressive decline in *Mllt3*, critical for the maintenance of the stem cell state^35^, mirrored by a slow increase in *Cdk6* that drives the exit from quiescence (or G0) and the G1-S transition in HSCs^36^ (Figure 1H). By contrast, the decline of the regulator of HSC self-renewal, *Mecom*^37^, took place abruptly at the ST-HSC/MPP border. A similarly abrupt decline was observed for *Hoxa9*, which coincided with the onset of lineage gene expression in pseudotime. Transcripts encoding respiratory enzymes (GO oxidative phosphorylation in Figure 1H) switched on when committed progenitors emerged, indicating reorganization of energy metabolism.

Taking chromatin accessibility and transcript data together, incremental and continuous gene-regulatory changes mark the transition from the HSC state to MPPs. These are followed by switch-like events, specifically a drop in both HOXA9 motif accessibility and transcript levels, as lineage commitment occurs. Among the known stemness regulators, HOXA9 showed the most distinctive chromatin accessibility footprint, with both motif accessibility and transcript levels being maintained along the HSC–MPP hierarchy and then dropping sharply upon lineage commitment, thereby distinguishing uncommitted from committed progenitors.

### Chromatin landscape indicates lineage restriction along multiple pathways

To characterize the rules of lineage specification, we asked which fate combinations, as evaluated by accessible chromatin, were present in progenitor cells. Using lineage probabilities (Figure 1C), we found that each mature cell fate emerged through a bifurcation of chromatin state in progenitors (Figure 2A). Progenitors assigned to the branch of a given lineage showed a marked increase in lineage probability after the bifurcation, whereas the remaining cells with low probability for the given lineage could still give rise to other lineages. Next, we computed successive lineage bifurcations to all individual cell fates (Figure 2B). This procedure assigns one or more of the six fates (Figure 1C) to each cell in the complete data set. Cells with all six fates correspond to stem cells, cells with two to five fates to oligolineage progenitors, and cells with one fate to unilineage progenitors. Of the 62 theoretically possible progenitor states (i.e., fate combinations), we found 60 in the data (Figure S3A shows the most frequent combinations). Of note, we recently indeed found experimental evidence for large numbers of fate combinations in progenitors based on barcoding experiments in native hematopoiesis (Cirovic et al., doi: https://doi.org/10.64898/2026.04.09.716798). However, the underlying ATAC-seq data are subject to experimental noise and this will likely cause spurious fate combinations (e.g., due to dropouts).

**Figure 2.**
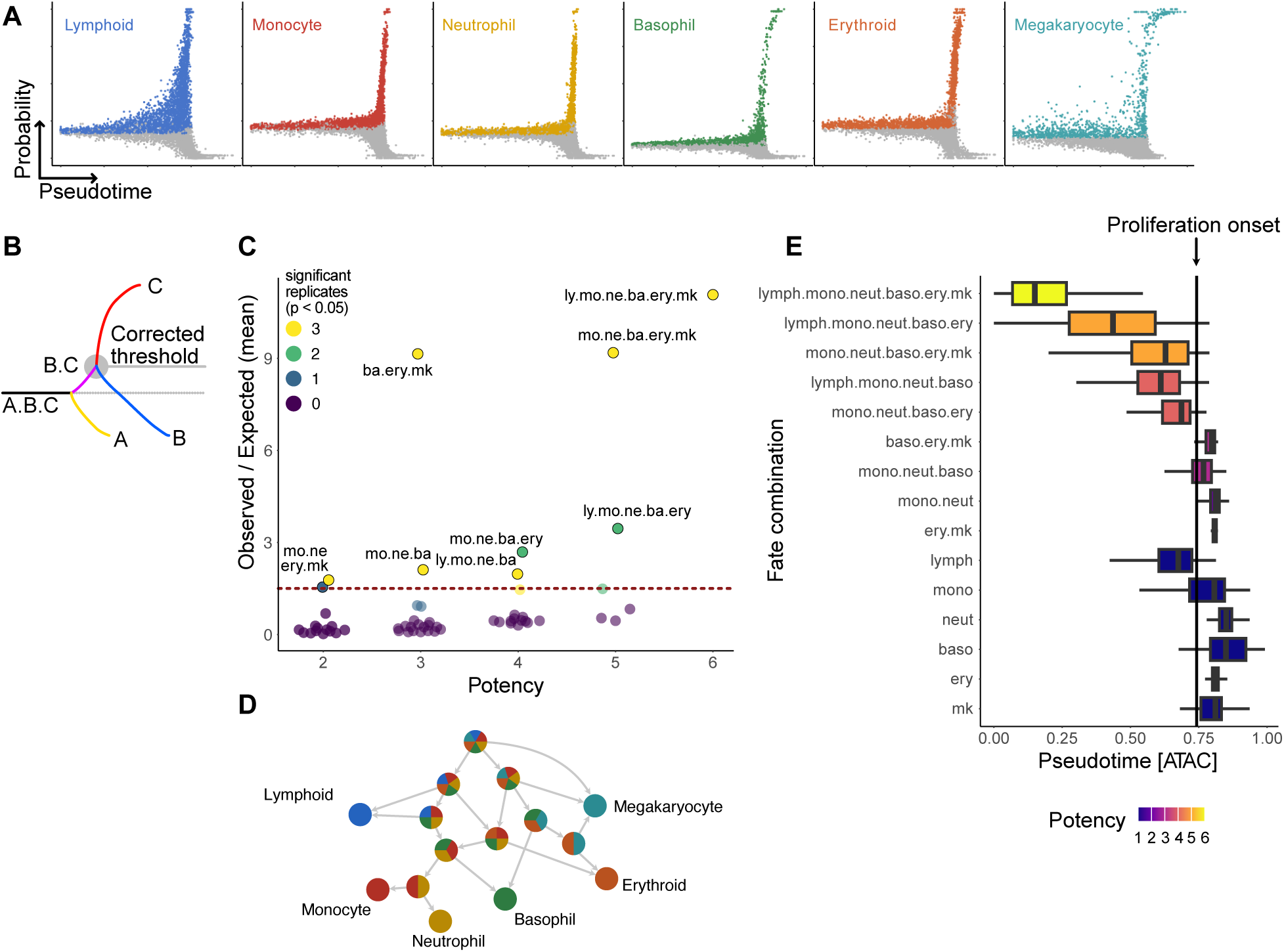
Chromatin state predicts hierarchy of oligo-lineage progenitors. (A) Single cells in pseudotime-probability scatterplots. Each scatter plot contains all cells that are arranged by pseudo-time (x-axis) and their inferred probability to commit to the indicated lineage (y-axis), using chromatin accessibility at hematopoietic enhancers. Cells are colored if they have the potential to commit to the respective lineage (Experiment 1). (B) Schematic representation of the iterative fate assignment model. Initially, cells whose lineage probability exceeds a data-derived threshold are assigned a potential fate. Further downstream, the threshold is recomputed to account for successive bifurcating events. (C) Oligo-lineage progenitors with particular combinations. We tested all fate combinations to determine whether they were enriched over occurrence by chance. The significance threshold for p = 0.05 (corrected for multiple testing) is indicated by the dashed line. Progenitors were included that were significantly enriched in at least one biological replicate (color code). Potency (x-axis) indicates the number of fates of a given progenitor. (D) Potential connections between the oligo-potent states detected in (C). (E) Pseudo-temporal distribution of progenitors with distinct combinations of possible fates. Only fate combinations that were statistically supported in at least one replicate are shown (see (C)). The black vertical line marks the time point at which half of the maximum proliferative activity is reached. Data are from Experiment 4; results for the other replicates in Figure S3.

To account systematically for the experimental noise, we assigned a progenitor state only when it occurred significantly more often in the noisy data than expected by chance. To this end, we developed an iterative version of the multi-set intersection test, the progenitor significance test (Methods). We first validated this approach computationally, generating synthetic data for two purely hypothetical differentiation landscapes: a traditional lineage tree containing only CMP, GMP, MEP, and CLP as progenitors (Figure S3B) and a cloud of all 62 possible progenitors, with all possible paths of step-wise lineage restriction (Figure S3C). When applying the progenitor significance test, we recovered exactly the progenitor types used to build the tree model (Figure S3D) and no significantly over-represented progenitor types for the cloud model (Figure S3E). Hence, for both sets of synthetic data, we correctly inferred the input progenitor structure. Next, we applied the progenitor significance test to the experimental data, yielding nine significant oligolineage progenitors (out of the original 60 fate combinations) (Figure 2C). This finding implies that the hematopoietic differentiation landscape is both more complex than the traditional lineage tree and more structured than a random progenitor cloud (Figure 2D).

In particular, we predicted two distinct tri-lineage progenitor populations that share basophil potential (mo.ne.ba and ba.ery.mk; Figure 2C), suggesting the existence of alternative differentiation routes (Figure 2D), as recently also shown for megakaryocytes^17,38^. Comparison of the pseudotime positions of cells assigned to the identified oligo- and unilineage populations showed that myeloid- or erythroid-restricted progenitors consistently emerged at pseudotime values after the point of proliferative activation of uncommitted progenitors (Figure 2E, Figure S3F).

### Uncommitted progenitors gain chromatin accessibility in S phase

To probe whether cell cycling and the emergence of differentiation-associated chromatin changes are mechanistically related, we first explored how genome-wide chromatin accessibility changes through the cell cycle.

As expected^26^, our data suggest that chromatin changes precede lineage gene induction (Figure S1). To detect differentiation-initiating chromatin changes, we identified transcriptionally uncommitted progenitors by the absence of lineage transcripts. Based on cell cycle phase-specific transcripts, we then defined uncommitted progenitors that are unambiguously in G1, S, or G2/M phase (Figure S4A). We reasoned that the DNA replication should leave a signature in the ATACseq data and, therefore, systematically searched for cell-cycle-associated chromatin accessibility features using unbiased machine learning (Figure S4B-E). We indeed found a strong cell-cycle signature in the ATAC-seq data from two out of the three biological replicates. In these two replicates, tagmentation was highly effective, as shown by the large DNA fragment diversity per cell (i.e. ∼38,000 unique fragments per cell for Exp1 and ∼46,000 for Exp4 at a mean sequencing depth of 100,000 fragments per cell), whereas the third replicate without a clear cell-cycle signature had lower fragment diversity (∼27,000 unique fragments for Exp3) (Figure S4F,G; Figure S5). In the two high-coverage replicates, we found that transposase insertion counts strongly increased over the cell cycle, roughly doubling from G1 to G2, consistent with replication (Figure 3A, Figure S6A).

**Figure 3.**
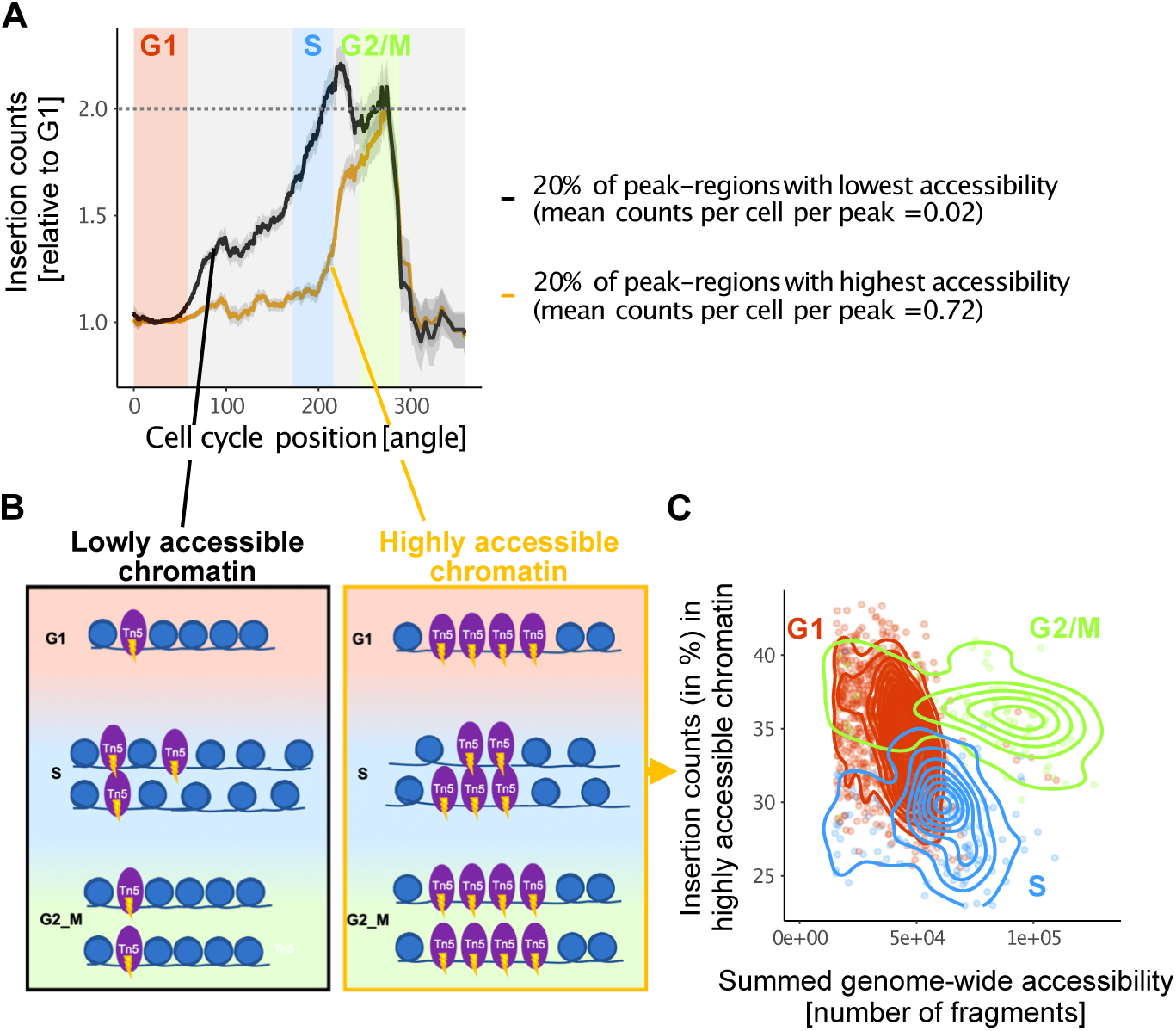
Cell-cycle signature in single-cell ATAC data. (A) Dynamics of chromatin accessibility (insertion counts) during cell cycle in transcriptionally uncommitted progenitors. Genomic regions with low G1 accessibility (black line) and high G1 accessibility (orange line) were selected and the factor by which the initial accessibility had changed was plotted. Values represent moving averages (window size of 15°) of cells that were ordered by cell-cycle position based on their RNA profiles (Figure S4A); positions of cells with unambiguous assignment of cell-cycle phase indicated by colors. Data from Experiment 4 (Experiment 1 in Figure S6A). (B) Cartoon of chromatin changes during cell-cycle progression. Blue circles represent nucleosomes with DNA (blue line). Purple ellipses represent dimers of the Tn5 enzyme used for ATACseq. Tn5 insertions (flashes) generate accessibility signals. At the level of individual fibers, chromatin generally becomes hyper-accessible after replication^54^ (left), whereas nucleosome-depleted regions become transiently more occupied with nucleosomes after replication and therefore show restricted accessibility for Tn5^39^ (right). (C) Two-dimensional plot separating cell-cycle phases by chromatin accessibility, using the total amount of accessible DNA per cell and the percentage of insertion counts in highly accessible chromatin as features. Cells are colored by cell-cycle phase according to their transcriptomes (see Figure S4A). Data from Experiment 4 (Experiment 1 in Figure S6B).

While all chromatin regions doubled the insertion count from G1 to G2, the kinetics of doubling depended on the initial state of the chromatin in G1. Chromatin regions that started with low G1 accessibility gained accessibility more rapidly during replication, relative to their state in G1, than regions with high starting accessibility in G1 (Figure 3A, Figure S6A). Mechanistically, the doubling of genome-wide accessibility in G2 (Figure 3A) is consistent G1-level accessibility being re-established on each chromatid (Figure 3B). Then, the transient increase by more than twofold at lowly accessible regions (Figure 3A, peak of the black curve in late S) indicates a genuine, transient rise in chromatin accessibility at individual fibers (Figure 3B, left). Conversely, the delayed increase in insertion counts at loci that were highly accessible in G1 phase is consistent with the previous finding that long stretches of nucleosome-depleted regions become transiently reoccupied by nucleosomes upon replication, before being remodeled to an open configuration (Figure 3B, right)^39^.

Thus far, we analyzed chromatin states averaged over the cell population of uncommitted progenitors to gain statistical power. Remarkably, however, the distinct dynamics of highly and lowly accessible chromatin loci through the cell cycle are visible also at the single-cell level (Figure 3C). Here, we plotted the insertion counts originating from highly accessible loci in G1 against the total accessibility per cell (measured by the number of sequenced unique DNA fragments per cell). These two features allow the separation of cells into different cell-cycle phases (with some overlap between G1 and S), with highly accessible loci trailing in S phase in this “phase plot” (Figure 3C, Figure S6B). Taken together, these findings reveal a prominent cell-cycle signature in high-coverage snATAC-seq data.

### Opening of lineage-associated chromatin regions at the onset of differentiation

We next asked which chromatin accessibility dynamics are associated with cell differentiation. To this end, we collected all chromatin loci that showed significant accessibility in one or more HSPC subpopulations (uncommitted progenitors, or progenitors showing specific lineage gene expression). First, we considered all regions with low accessibility in uncommitted progenitors (the 20% of peak regions with lowest G1 accessibility analyzed in Figure 3A) and asked whether they opened during myeloid differentiation, or megakaryocyte-erythroid differentiation, or both. The vast majority of closed chromatin loci in uncommitted progenitors (>80%) opened during one or both differentiation processes, with myeloid differentiation showing the most pronounced opening (Figure 4A, black bars). By contrast, loci that were already open in uncommitted progenitors (the 20% of peak regions with highest G1 accessibility analyzed in Figure 3A) mostly did not open further (Figure 4A, orange bars). Thus, the onset of differentiation is linked with extensive opening of chromatin regions that were closed in uncommitted progenitors. Next, we asked whether open chromatin regions in uncommitted progenitors closed at the onset of differentiation. The majority of open regions stayed open upon onset of differentiation and did not close (Figure 4B, orange bars). Taken together, myeloid and megakaryocyte-erythroid differentiation is associated predominantly with the opening of chromatin loci that were closed in uncommitted progenitors.

**Figure 4.**
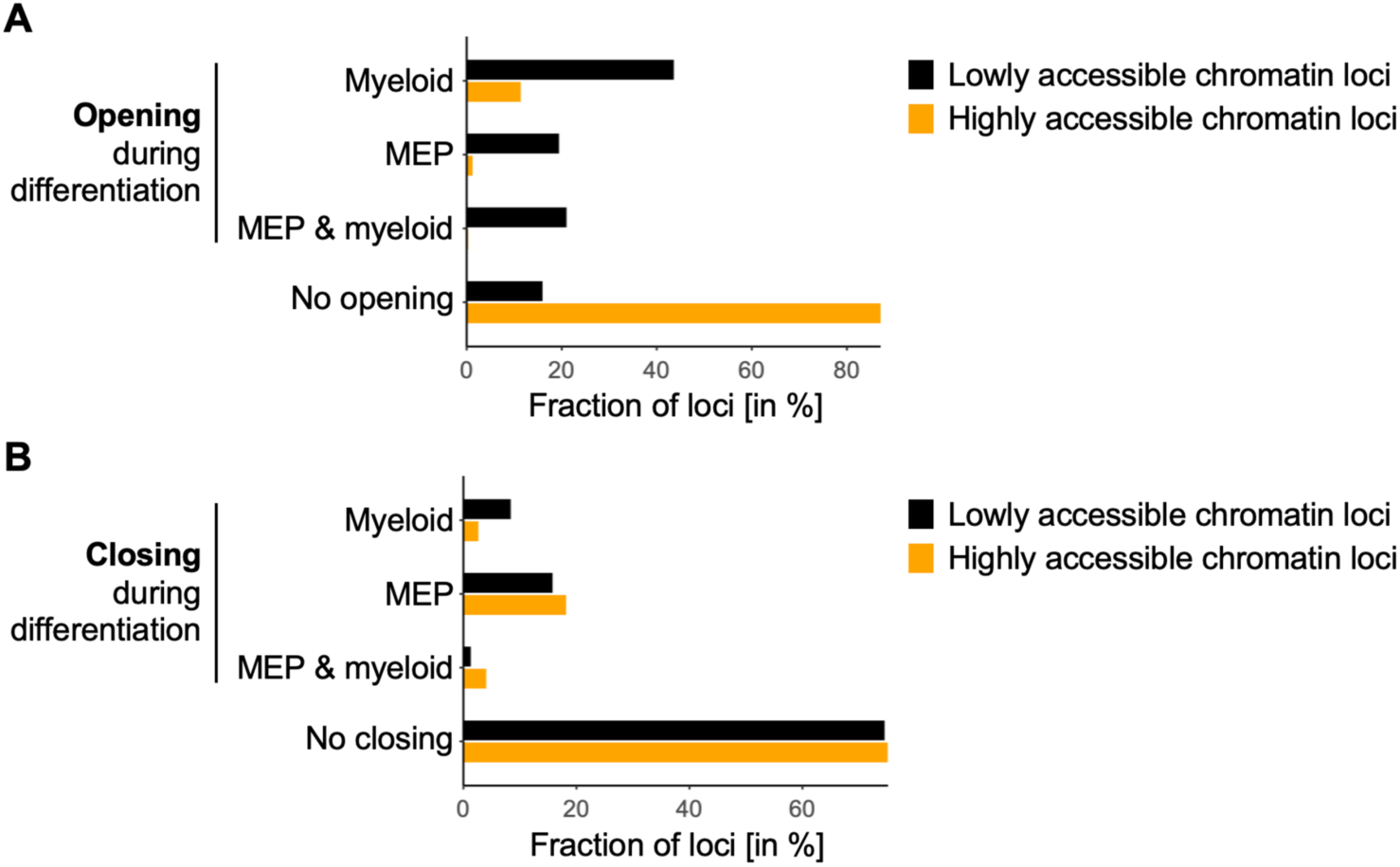
Myeloid commitment is marked by opening of previously lowly accessible loci. (A) Extent of chromatin opening during differentiation. Black bars, chromatin regions that were lowly accessible in uncommitted HSPCs; Orange bars, regions that were already highly accessible before lineage commitment. (B) Extent of chromatin closing during differentiation. Black bars, chromatin regions that were lowly accessible in uncommitted HSPCs; Orange bars, regions that were already highly accessible before lineage commitment. Data from the biological replicate with highest genome coverage (Experiment 4).

### Myeloid regulatory loci open during S phase

Our above finding of opening of lineage-specific loci (cf. Figure 4A) was based on pseudo-temporal ordering of cells along lineage trajectories. To probe chromatin opening events at finer resolution in relation to the cell cycle, we compared uncommitted progenitors in different cell-cycle phases, as follows: We identified, within transcriptionally undifferentiated cells, cell pairs with similar transcriptomes where one cell was in G1 and the other in either S or G2/M, which we refer to as transcriptionally matched neighbors, or simply matched neighbors (Figure 5A). We selected on average 248 of matched neighbors per replicate (124 cells in S; 124 in G1). First, we validated that matched neighbors were equally undifferentiated (Figure S7). Specifically, all cells had high expression of *Hoxa9* and basal expression of *Spi1* (coding for PU.1), characteristic of the uncommitted progenitor state^40–43^ (Figure 5B). Moreover, cells expressed low levels of the myeloid master regulator *Cebpa*^44,45^ (Figure 5B) and the erythro-megakaryocytic master regulator *Gata1*^46^, further confirming their undifferentiated state (Figure S8).

**Figure 5.**
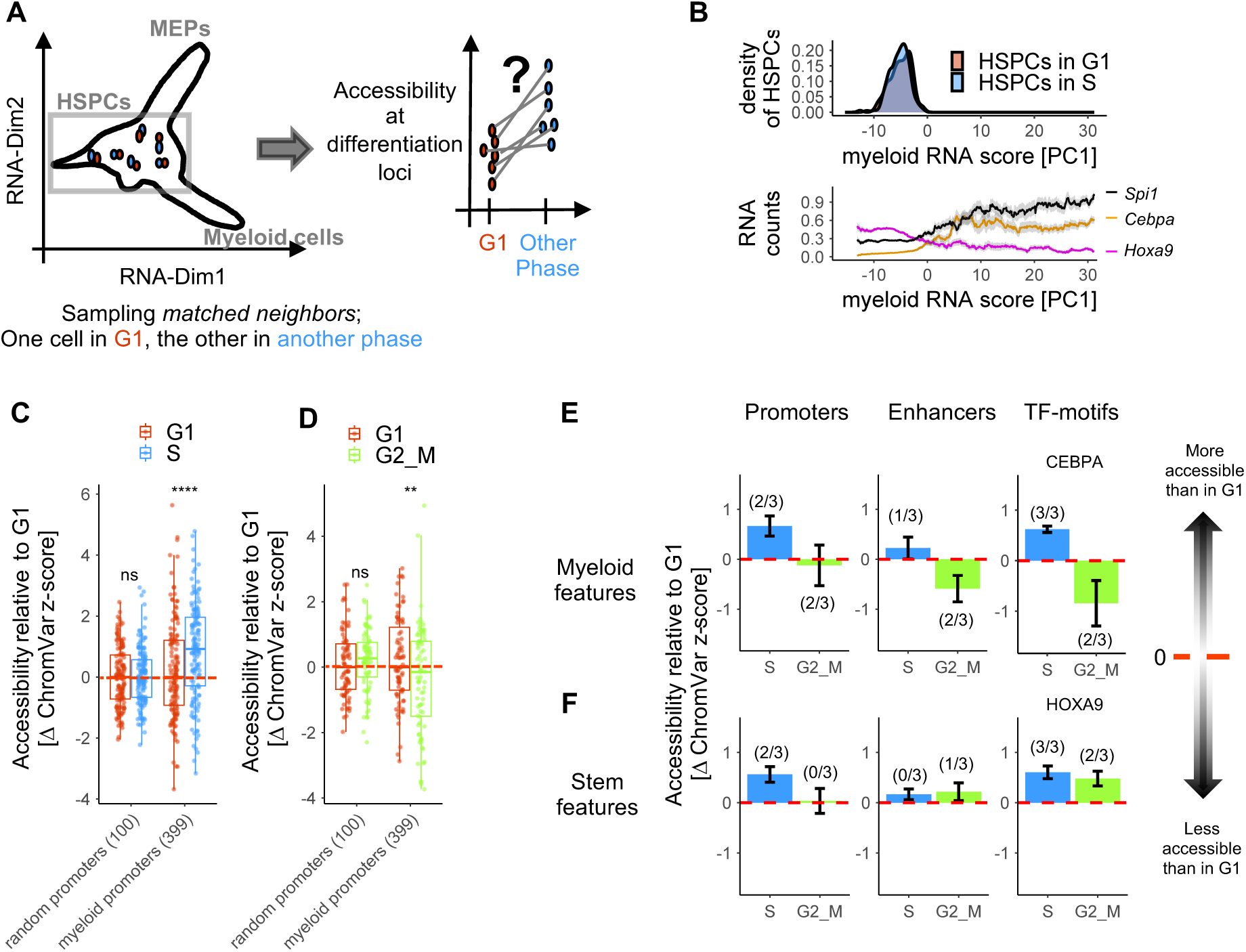
During S phase, uncommitted HSPCs open myeloid regulatory loci. (A) Illustration of the cell selection used to compare chromatin accessibility between transcriptionally matched cells in different cell-cycle phases. Selected HSPC pairs are transcriptionally uncommitted. (B) Position of transcriptionally matched HSPCs in G1 and S phase along the RNA-based differentiation trajectory identified by PCA (PC1), ranging from transcriptionally uncommitted cells (PC1 < 0) to myeloid-committed cells (PC1 > 0). Colored lines represent moving averages of RNA counts for key transcription factors associated with multipotency (*Hoxa9*) and myeloid differentiation (*Spi1* and *Cebpa*). Grey ribbons indicate the standard error of the mean. Data shown are from Experiment 4. (C) Normalized chromatin accessibility of uncommitted G1-phase cells (red dots) and transcriptionally matched cells in S phase (blue dots). Left boxplot, accessibility at 100 randomly selected promoters, each dot representing one cell. Normalization results in indistinguishable z scores. Right boxplot, accessibility at 399 promoters that open during myeloid lineage differentiation, each dot representing one cell. P-values were calculated using a paired Wilcoxon test (indicated significance levels: p<0.0001: ****; p<0.001: ***; p<0.01: **; p<0.05*). Data shown are from Experiment 4. For plotting accessibility relative to G1 cells, ChromVar z-scores were aligned to place the median of G1 cells at zero. (D) Comparison of normalized chromatin accessibility between uncommitted G1-phase cells (red dots) and transcriptionally matched cells in G2/M phase (green dots), as described in C. (E) Cell-cycle-associated chromatin changes at myeloid promoters, enhancers and CEPBA transcription factor (TF) motifs, in S phase relative to G1 (blue bars) and G2/M phase relative to G1 (green bars). Each bar represents the mean difference of cells in the respective phase to transcriptionally matched cells in G1 aggregated across all three replicates. Error bars indicate the standard error of the mean. Numbers above each bar indicate in how many of the three mice the deviation from G1 (red dashed line) was statistically significant (paired Wilcoxon test, *p* < 0.05). For data of individual mice see Figure S9. (F) Cell-cycle-associated chromatin changes at promoters, enhancers and HOXA9 transcription factor (TF) motifs associated with the uncommitted HSPC state, in S phase relative to G1 (blue bars) and G2/M phase relative to G1 (green bars). Details as in E.

We next examined accessibility of differentiation-associated chromatin loci in single cells, while controlling for baseline accessibility using ChromVar^47^ (cf. Figure 3A). When choosing 100 promoters at random, ChromVar corrected for generic cell-cycle effects unrelated to differentiation, as seen by matching ChromVar-derived z-scores for cells in G1 and other cell-cycle phases (Figure 5C,D, left in both). This normalization accounts for the generic behavior of chromatin regions during the cell cycle, shown in Figure 3A. In stark contrast to this baseline, we found that promoters of myeloid lineage genes showed higher accessibility during S phase than expected (Figure 5C, right). This S phase-linked opening of myeloid promoters was detected in all three biological replicates and was significant in two of them (Figure 5E, left, the bars show the average signal across all replicates, while the numbers in brackets show in how many replicates the change was significant; see also Figure S9A for the individual replicates). Moreover, the binding sites of the myeloid master regulator, CEBPA, strongly gained accessibility in S phase across all replicates (Figure 5E, right), whereas the S phase-specific opening signal for myeloid enhancers was weaker (Figure 5E, middle). These data imply that the opening of myeloid promoters (as part of the chromatin regions assayed in Figure 4A) begins in S phase.

In G2 phase cells, remarkably, myeloid-associated regions (promoters, enhancers and CEBPA motifs) tended to be less accessible than in G1 phase cells (Figure 5E; Figure S9A). This observation is consistent with a single cell cycle in uncommitted progenitors being able to initiate myeloid differentiation. Undifferentiated progenitors in both G1 and G2 phases will then be in a state prior to myeloid commitment, whereas in a subset of undifferentiated S phase cells, myeloid differentiation will be initiated.

We also found that stem-cell-specific regulatory chromatin – promoters of stemness genes and HOXA9 binding motifs – had a specific opening signal in S phase (Figure 5F). In contrast to myeloid regions, this signal persisted into G2. This finding suggests that S phase transition could alternatively reinforce the existing uncommitted progenitor state and, moreover, that maintaining the undifferentiated progenitor state and initiating myeloid differentiation could be competing processes.

### S-phase opening of myeloid loci is enhanced in emergency myelopoiesis

To further probe the relation between S phase transition and onset of myeloid differentiation, we acutely enhanced myelopoiesis by systemic infection^48^. In this process, we found previously that the proliferation of uncommitted progenitors is strongly accelerated at 24 hours after infection, followed by recovery to baseline proliferation by day 3^32^. To investigate whether accelerated proliferation and enhanced myelopoiesis are coordinated, we induced sepsis by intraperitoneal injection of cecal slurry and isolated HSPCs after 24 hours. We performed three replicate sepsis experiments (each done in parallel with one of the steady-state experiments analyzed thus far), and indeed observed the hallmarks of a systemic inflammatory response (Figure S10). Specifically, we harvested HSCs, ST-HSCs, MPPs, and LK cells from the bone marrow and generated single-cell multiome data, resulting in a total of 21,811 high-quality cells for joint analysis of chromatin accessibility and transcriptome (3,626 HSCs, 817 ST-HSCs, 7,646 MPPs and 9,722 LK cells, from three replicate experiments).

To integrate these data, we projected the sepsis ATAC-seq data into the chromatin state space of steady-state hematopoiesis and found the same major lineages in the sepsis response as previously detected in steady-state (Figure 6A,B; Figure S11A,E). Sepsis caused a shift of cell densities toward more differentiated cell states, consistent with a wave of enhanced differentiation (Figure 6C; Figure S11B,F). In particular, we detected increased probability of myeloid differentiation in LSK cells (Figure 6D; Figure S11C,G), which agrees with the transiently enhanced production rate of neutrophil granulocytes that we observed previously by time-resolved fate mapping^33^. The proliferative activation of HSCPs was accelerated, yielding an earlier proliferation onset in pseudotime (defined as 50% of the maximal fraction of S/G2 cells observed) (Figure 6E, upper row). The 50% points of lineage potentials now trailed the proliferation onset for all lineages (including the lymphoid lineage), showing that in sepsis proliferative activation preceded all lineage decisions. Remarkably, however, the onset points of differentiation were hardly shifted from their respective steady-state counterparts (Figure 6E; Figure S11D,H). Taken together, these findings suggest that the underlying progression of enhancer states from HSCs to MPPs and then to different lineage pathways was very similar in steady state and in emergency myelopoiesis, whereas the flux of myeloid cell production was increased.

**Figure 6.**
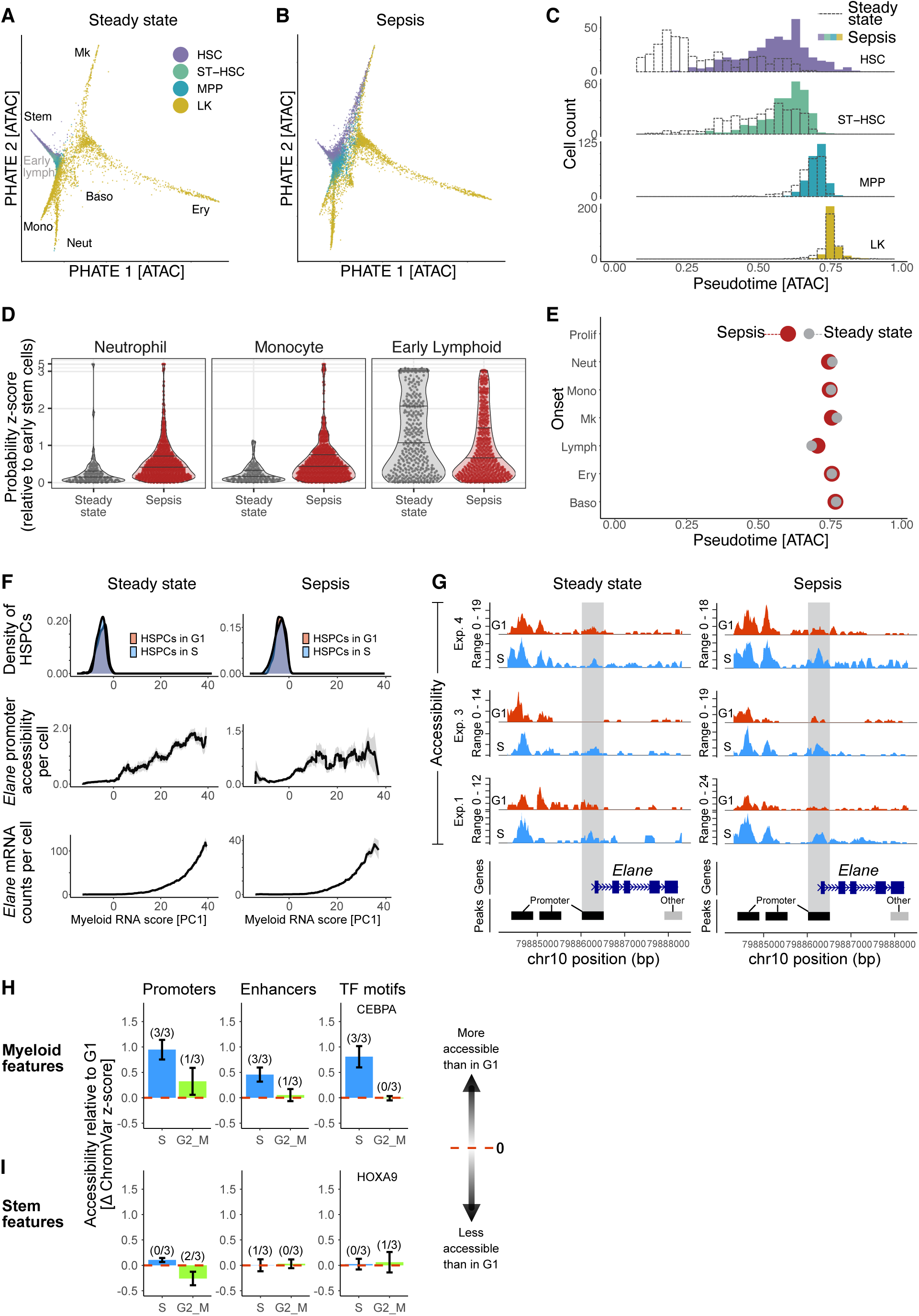
Sepsis enhances S phase-specific opening of myeloid regulatory loci. (A, B) PHATE visualization of HSPCs under steady-state conditions (A) and in response to sepsis (B). Cells from the sepsis condition were projected onto the control dataset. Cells are colored according to FACS-sorted populations (HSC, ST-HSC, MPP, LK) and branches are labelled according to lineage endpoints. As in Figure 1B, accessibilities of hematopoietic enhancers were used as features. (C) Density distribution of sorted cell populations (HSC, ST-HSC, MPP, LK) along pseudotime, comparing control (dotted lines) and sepsis conditions (filled histograms). (D) Distributions of lineage probability scores in LSK cells of control and sepsis conditions for the indicated lineages. Z-scores were computed by subtracting the mean probability observed in early stem cells and dividing by the corresponding standard deviation. Scores < 0 are not shown. (E) Onset of cell cycle activation (top row) and lineage differentiation (below) during sepsis (red dots), compared to steady state (grey dots). Onset was computed as half-maximum constant for the monotonic splines fitted to the non-G1 fraction (top row) or the lineage probabilities. (F) Accessibility and transcription dynamics at the *Elane* locus during myeloid differentiation in steady state (left panels) and during sepsis (right panels). The x-axis defines the RNA-based differentiation trajectory, from transcriptionally uncommitted cells (PC1 < 0) to myeloid-committed cells (PC1 > 0) (as in Figure 5B). Top row, Cell-density curves mark the positions of uncommitted, transcriptionally matched G1 and S phase cells; only uncommitted cells were used for analyses from (F) to (I). Middle row, Promoter accessibility reflects insertion counts within the 500 bp wide promoter peak overlapping the TSS of *Elane*. Bottom row, Average number of *Elane* transcripts detected per cell. All values are centered moving averages (window size = 4 units), based on data from the steady state and sepsis mouse of Experiment 4. Gray ribbons denote the standard error of the mean. (G) Chromatin accessibility at the *Elane* locus. Centered moving sum of chromatin accessibility (Tn5 insertion counts) at the *Elane* locus (moving window size = 100 bp). Tracks depict data from transcriptionally matched, uncommitted cells in G1 (red) and S (blue) phases for untreated (left) and septic (right) mice. Each marked peak spans 500 bp. Gray shading indicates the promoter peak overlapping the transcription start site (TSS). (E) Cell-cycle-associated chromatin changes at myeloid promoters, enhancers and CEPBA transcription factor (TF) motifs, in S phase relative to G1 (blue bars) and G2/M phase relative to G1 (green bars) in sepsis. Each bar represents the mean difference of cells in the respective phase to transcriptionally matched cells in G1 aggregated across all three replicates. Error bars indicate the standard error of the mean. Numbers above each bar indicate in how many of the three mice the deviation from G1 (red dashed line) was statistically significant (paired Wilcoxon test, *p* < 0.05). For data of individual mice see Figure S9. (F) Cell-cycle-associated chromatin changes at promoters, enhancers and HOXA9 transcription factor (TF) motifs associated with the uncommitted HSPC state, in S phase relative to G1 (blue bars) and G2/M phase relative to G1 (green bars). Details as in E. (A-E) Analyses based on the control and sepsis mouse from Experiment 1. See Figure S11 for other replicates.

Next, we assessed the extent of S-phase-linked chromatin opening at individual myeloid differentiation-associated loci in sepsis. We focused on the *Elane* gene locus, whose transcriptional activation is an early marker of neutrophil commitment^49,50^ and is regulated through the binding of the myeloid transcription factors (TFs) CEBPA and PU.1 to its promoter^51^. Importantly, *Elane* is not expressed in uncommitted progenitors (in which we analyze chromatin opening) but becomes strongly transcribed downstream (Figure 6F, top and bottom). Notably, opening of the *Elane* promoter preceded transcription (Figure 6F, middle). While in steady-state hematopoiesis, the signal for selective opening of the *Elane* promoter in S phase was comparatively weak (Figure 6G, left), we observed a clear S phase-specific accessibility gain in all three sepsis replicates (Figure 6G, right). We found a similar behavior of the accessibility of the *Mpo* proximal enhancers (Figure S12). These data on the *Elane* and *Mpo* loci suggest that the S phase-coupled opening of myeloid loci in uncommitted progenitors is augmented in emergency myelopoiesis.

When quantifying chromatin accessibility across myeloid promoters, enhancers, or CEBPA binding sites (as done above for steady state), we detected pronounced and significant S phase-coupled chromatin opening at all three types of myeloid regulatory loci in all sepsis experiments (Figure 6H, Figure S9B for values of individual biological replicates). This stood in sharp contrast to promoters and enhancers of stemness genes, as well as HOXA9 binding motifs, which did not gain accessibility during S phase (Figure 6I). We therefore asked if differences in chromatin opening at TF binding motifs were a consequence of altered TF expression in sepsis. To assess this possibility, we quantified transcripts of *Hoxa9*, as well as baseline transcripts of *Cebpa* and *Spi1* in uncommitted progenitors in steady state and during sepsis. All three transcription factors had very low counts, precluding robust detection of differences. However, in the experiment with the highest RNA coverage, we detected a small but significant decrease of *Hoxa9* expression in sepsis (Figure S13), which raises the possibility that *Hoxa9* expression could be quantitatively reduced in uncommitted progenitors in sepsis. Taken together, in emergency myelopoiesis transcriptionally uncommitted progenitors show strong opening of myeloid loci in S phase (promoters, enhancers and CEBPA binding sites) and no opposing opening of stemness-associated loci.

### HOXA9 motif accessibility predicts opening of myeloid regulatory regions in S phase

In uncommitted progenitors in steady-state hematopoiesis, both stemness-associated regulatory regions and myeloid regulatory regions gained accessibility during S phase (Figure 5E,F), while in emergency myelopoiesis, the opening of stemness-associated regulatory was greatly diminished (Figure 6H,I). These observations raise the question whether chromatin changes that reinforce the uncommitted state, on one hand, and changes that initiate myeloid differentiation, on the other hand, are mutually exclusive in individual cells or co-occur. To answer this question with appropriate statistical power, we focused on subpopulations of cells instead of single cells per se. We computationally sorted subpopulations of uncommitted progenitors in S phase with low or high HOXA9 motif accessibility (schematic in Figure 7A, top, choosing specifically the lower quartile as HOXA9-low cells, and the upper quartile as HOXA9-high cells). We then probed for S phase-specific opening of particular chromatin regions separately in HOXA9-low and HOXA9-high cells.

**Figure 7.**
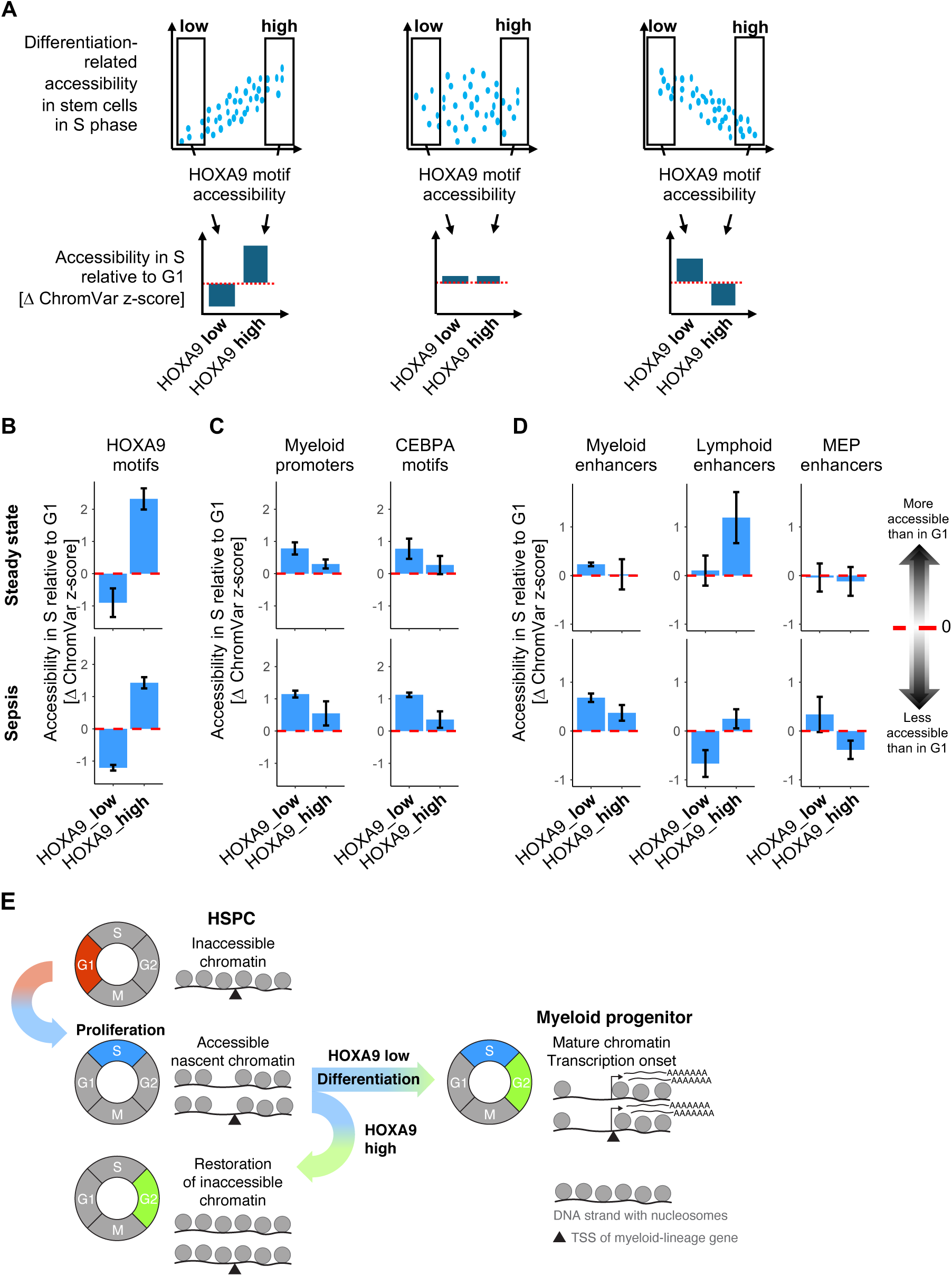
Chromatin occupancy by HOXA9 in uncommitted progenitors predicts whether S phase will initiate myelopoiesis. (A) (Top row) Schematic of possible relations between the accessibility at HOXA9 sequence motifs and the accessibility at differentiation-associated loci, quantified as before in uncommitted progenitors in S phase. (B-D) Accessibility relative to G1 phase at distinct loci in uncommitted S phase cells, separated according to HOXA9 motif accessibility (lower 25%, HOXA9-low; upper 25%, HOXA9-high). (B) Accessibility at HOXA9 binding motifs, confirming the computational cell sort. (C) Accessibility at myeloid promoters (left panels) and at CEBPA binding motifs (right panels). (D) Accessibility at lineage-specific enhancers, for myeloid, lymphoid and megakaryocyte-erythroid (MEP) differentiation. (E) Model for initiation of myeloid differentiation. Uncommitted HSPCs generally exhibit inaccessible chromatin at myeloid-regulatory regions. During S phase these loci become transiently more accessible. Cells with low HOXA9 motif occupancy show particularly large accessibility gains and may commit to myeloid differentiation, whereas cells with high HOXA9 motif occupancy re-establish a condensed chromatin state at myeloid loci.

The HOXA9-high population showed an S phase-specific increase in HOXA9 motif accessibility (Figure 7B), consistent with the reinforcement of the uncommitted state observed above for unsorted uncommitted progenitors (cf. Figure 5F). This occurred most strongly in steady state, and to a lesser extent in sepsis. By contrast, the HOXA9-low population did not show this effect (Figure 7B). These data suggest that HOXA9-high and HOXA9-low uncommitted progenitors are functionally distinct subpopulations. Indeed, the HOXA9-low progenitors, but not the HOXA9-high progenitors, strongly gained accessibility at myeloid regulatory loci during S phase (Figure 7C). This anti-correlation between HOXA9 motif accessibility and opening of myeloid regulatory loci was conserved across steady state and emergency myelopoiesis, and was more pronounced in the latter (Figure 7C). Myeloid opening coincides with the closing of HOXA9 motifs in emergency myelopoiesis in HOXA9-low progenitors (Figure 7B). These findings indicate that processes which safeguard the chromatin profile of uncommitted progenitors compete with processes driving the transition of the chromatin state to myeloid differentiation during S phase.

In line with the putative role of HOXA9 to prevent excessive opening of myeloid regulatory regions during S phase, it has been reported that HOXA9 supports the preservation of uncommitted multipotent progenitors with lymphoid differentiation potential^42,52^. Indeed, we found that high HOXA9 motif accessibility coincides with increased accessibility at lymphoid enhancers in S phase (Figure 7D, top). In emergency myelopoiesis, however, accessibility of lymphoid enhancers is hardly increased in HOXA9-high S phase cells and reduced in HOXA9-low S phase cells (Figure 7D, bottom), which is consistent with the observed transient suppression of lymphopoiesis in sepsis^33^. Opposed to the behavior of lymphoid enhancers, myeloid enhancers selectively open in sepsis, primarily in HOXA9-low cells. Interestingly, we observed only minor changes in accessibility at MEP enhancers. In summary, we found that the reinforcement of the uncommitted state, marked by high HOXA9 motif accessibility, and the initiation of myeloid differentiation are mutually exclusive events in multipotent progenitors.

## DISCUSSION

In this study, we generated a comprehensive single-cell multiome data set for steady-state and emergency hematopoiesis that allowed us to probe the onset of lineage differentiation in uncommitted progenitor cells. We detect coherent opening of lineage-associated chromatin loci in MPPs that are transcriptionally uncommitted (i.e., do not show elevated transcript levels of lineage-specifying transcription factors). This finding argues against the long-standing idea that random fluctuations in the expression of lineage-specifying transcription factors initiate stem cell differentiation. Of note, a prior time-lapse imaging study of PU.1 and GATA-1 dynamics in HSPCs ex vivo failed to find evidence for random fluctuations in either protein driving lineage decisions^53^, in line with our finding in the intact organism. Instead, we uncovered a link between the proliferative activation occurring during the differentiation of hematopoietic stem cells and the onset of lineage differentiation itself. We found that proliferation accelerated in uncommitted progenitors before lineage-specific chromatin loci opened, both in steady state and in emergency myelopoiesis. Subsequently, chromatin dynamics in the S phase played a pivotal role for myeloid lineage decisions.

Based on these findings lead, we propose a differentiation model in which commitment to the myeloid lineage is initiated during S phase in rapidly proliferating progenitors (Figure 7E). Multipotent progenitors progressing through DNA replication can either preserve condensed chromatin on nascent DNA at regions regulating myeloid transcription—thereby maintaining their multipotency—or lose this chromatin structure and initiate the transcription of lineage-specific genes. Since the latter scenario is predominantly observed in cells with reduced accessibility at HOXA9 motifs, this factor (and possibly other stemness regulators) may promote a replication mode that ensures faithful transmission of the epigenetic landscape to both nascent chromatin fibers, thereby supporting self-renewing divisions of uncommitted progenitors. Loss of this self-renewing replication mode, possibly upon loss of the respective niche signals, will then trigger progenitor differentiation as a default program.

Two recent studies in cell culture are consistent with synthesis of (nascent) chromatin facilitating *de novo* gain of chromatin accessibility. First, in mammalian cells, nascent DNA is generally wrapped less tightly around nucleosomes and, after replication, this loose chromatin structure requires several hours to return to the steady-state configuration^54^. Second, H3K27me3 histone marks, which mediate the condensed chromatin structure at inactive promoters, are transiently diluted for up to several hours after replication in cultured human and murine HSPCs^23^.

Furthermore, Petruk et al.^23^ reported that cytokine-driven myeloid and erythroid differentiation of human HSPCs relies on binding of lineage-TFs, such as CEBPA, to nascent DNA. Thus, our *in vivo* finding that CEPBA motifs open specifically in S phase in uncommitted progenitors strongly suggests that these mechanisms initiate myeloid lineage choice during native hematopoiesis. This is further supported by the concomitant opening of myeloid promoters and enhancers.

Our observation that pronounced S phase-coupled opening of myeloid regulatory regions occurs only in a fraction of uncommitted progenitors suggests the existence of distinct DNA replication conditions that either preserve epigenetic repression of lineage genes or permit the opening of their regulatory elements. Related to this idea, Do et al.^24^ found in a CRISPR screen that diverse perturbations that stressed replication also promoted the onset of cell differentiation in S phase in cultured hematopoietic progenitors. In line with this, we found that increased proliferation of progenitors *in vivo* during sepsis, and thus likely elevated replicative stress, coincided with enhanced opening of myeloid regulatory regions in uncommitted progenitors. Furthermore, the co-occurrence of reduced HOXA9 motif accessibility in uncommitted HSPCs and accessibility gains at myeloid regulatory regions might indicate elevated replication stress in those cells, both in steady-state and emergency hematopoiesis. Indeed, Lynch et al.^55^ reported that HOXA9 and β-catenin modulate DNA replication in HSPCs in a compensatory manner. They find that HOXA9 and β-catenin largely bind to the same genomic regions and jointly support efficient fork progression and maintenance of genomic integrity. Adding on the critical role of replication fork properties, an *in vivo* study demonstrated that the association of PRC1 with the replication fork via PCGF1 is required to prevent premature myeloid differentiation of HSPCs^56^. Together with our results, these reports suggest that DNA replication conditions can tip the balance from self-renewal to onset of myeloid differentiation.

Furthermore, our findings point to the possibility that HOXA9 not only enables self-renewing divisions of hematopoietic progenitors^55,57,58^ by limiting S phase-linked activation of myeloid regulatory loci (Figure 7C) but also supports lymphoid enhancer opening (Figure 7D), thus preserving the lymphoid potential of uncommitted progenitors. Lymphoid enhancer opening in S phase cells with high HOXA9 motif accessibility is in line with the known direct transcriptional activation of *Flt3* by HOXA9^42^. *Flt3* expression marks lymphoid-primed progenitors that are fully multipotent^59,60^.

Importantly, our study introduces a mechanism for the initiation of myeloid differentiation that operates via the licensing of chromatin structure for the action of myeloid transcription factors. This model explains the puzzling observation that myeloid transcription factors PU.1 and CEBPA are already expressed at low levels in stem cells without inducing myeloid differentiation or antagonizing erythro-megakaryocyte fate^53^. This will be the case because corresponding target regions – promoters and enhancers – remain largely inaccessible to these transcription factors in stem cells, while chromatin opening in rapidly cycling progenitors allows CEBPA binding that, in turn, will trigger auto-activating and cross-repressing transcriptional programs.

### Limitations of the study

While our approach revealed that transcriptionally uncommitted HSPCs acquire accessibility at myeloid regulatory loci during S phase, it does not provide information about the fate of these cells. However, both the pseudotime ordering and the enhanced S phase opening of myeloid loci during sepsis, a condition characterized by increased myelopoiesis, suggest that S phase-coupled chromatin opening may indeed be followed by myeloid differentiation. Barcoding of single HSPCs in situ may further help resolve lineage fates, specifically, the progenitor types that were computationally predicted. Furthermore, while our analysis revealed that myeloid differentiation-associated loci become significantly more accessible during S phase, it cannot resolve whether this opening occurs exclusively behind the replication fork at the newly assembled chromatin fiber, or also at the parental fiber. Based on the reported properties of nascent chromatin^23,54^, it is likely that the enhanced accessibility signal in S phase comes from newly assembled chromatin.

## Supplementary Figures

**Figure S1.**
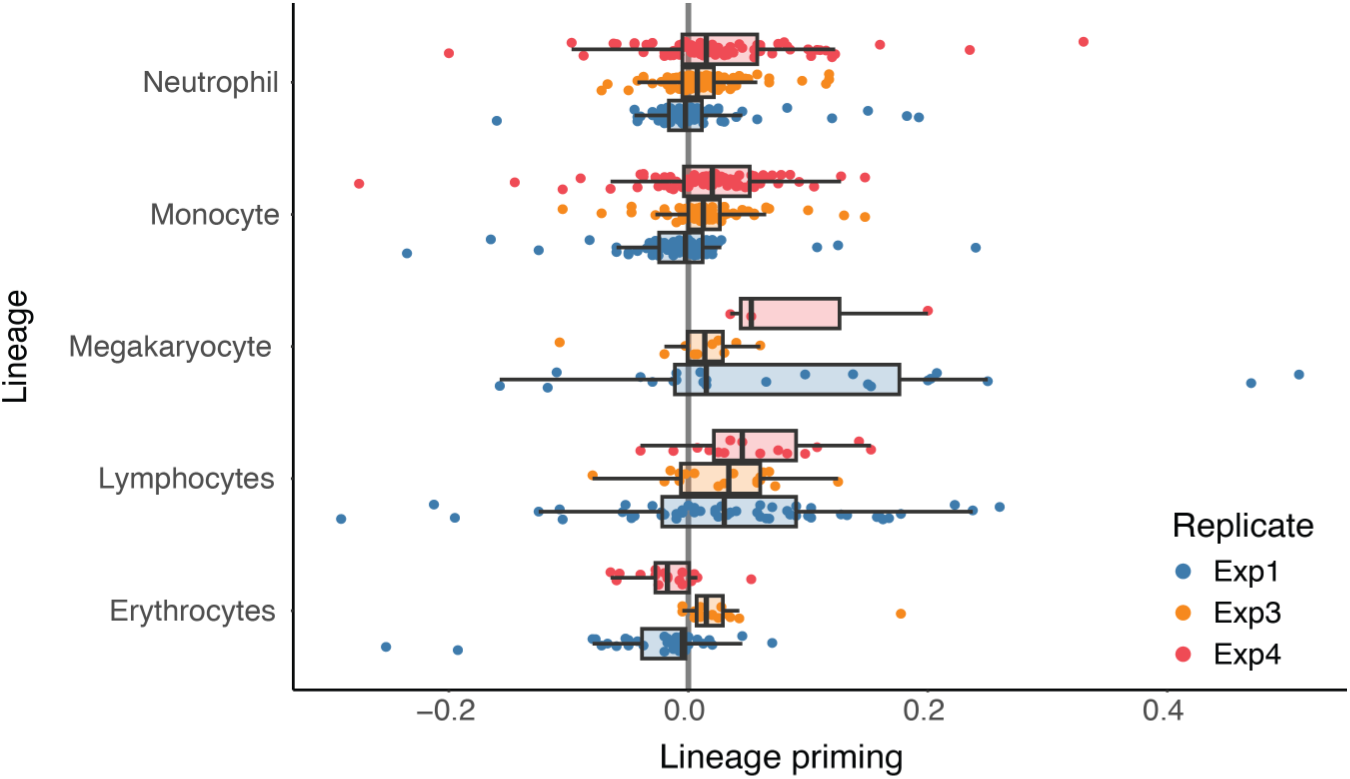
Chromatin opening precedes transcription. Lineage priming detected across replicates and lineages. Priming is measured as the difference between the half-maximum point of monotonic splines fitted to pseudotemporal trends of lineage-specific markers across RNA features (normalized expression) and the opening of the TSS+gene body measured by ATAC features (Gene Score Matrix).

**Figure S2.**
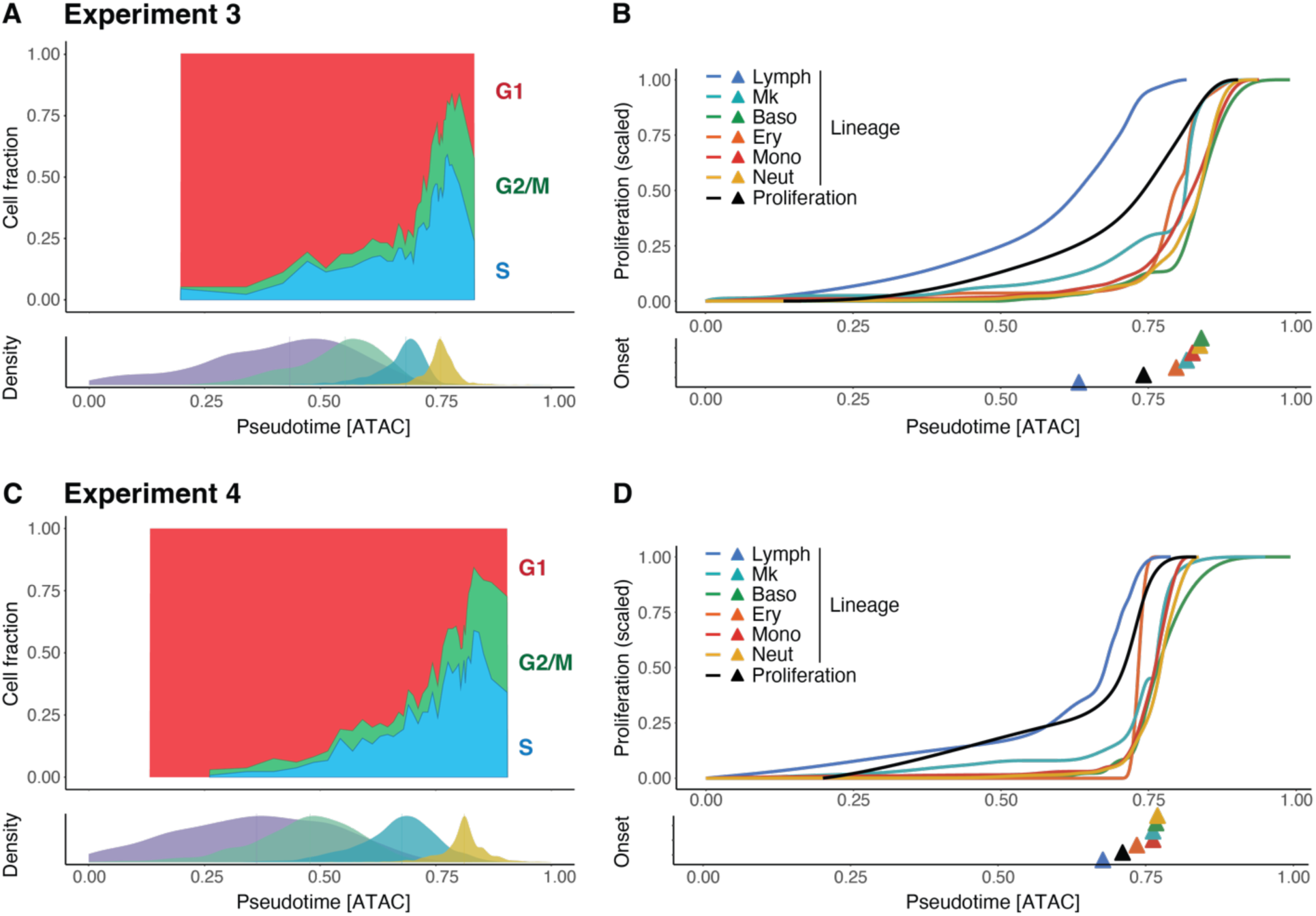
Proliferative activation precedes hematopoietic lineage specification. (A,B) Results from the untreated mouse of experiment 3. (A) (Top) Moving average of the fraction of cells in G1, S and G2/M phases of the cell cycle along differentiation pseudotime. (Bottom) Distributions of HSCs, STs, MPPs and LK cells (from left to right) along pseudotime. (B) (Top) Colored lines: fate probability inferred by CellRank as a function of pseudotime. Black line: Moving average of the fraction of cells that are not in G1 phase. (Bottom) Each triangle marks the point in pseudotime at which the corresponding line reaches the half of its maximum value. (C, D) Results from the untreated mouse of experiment 4.

**Figure S3.**
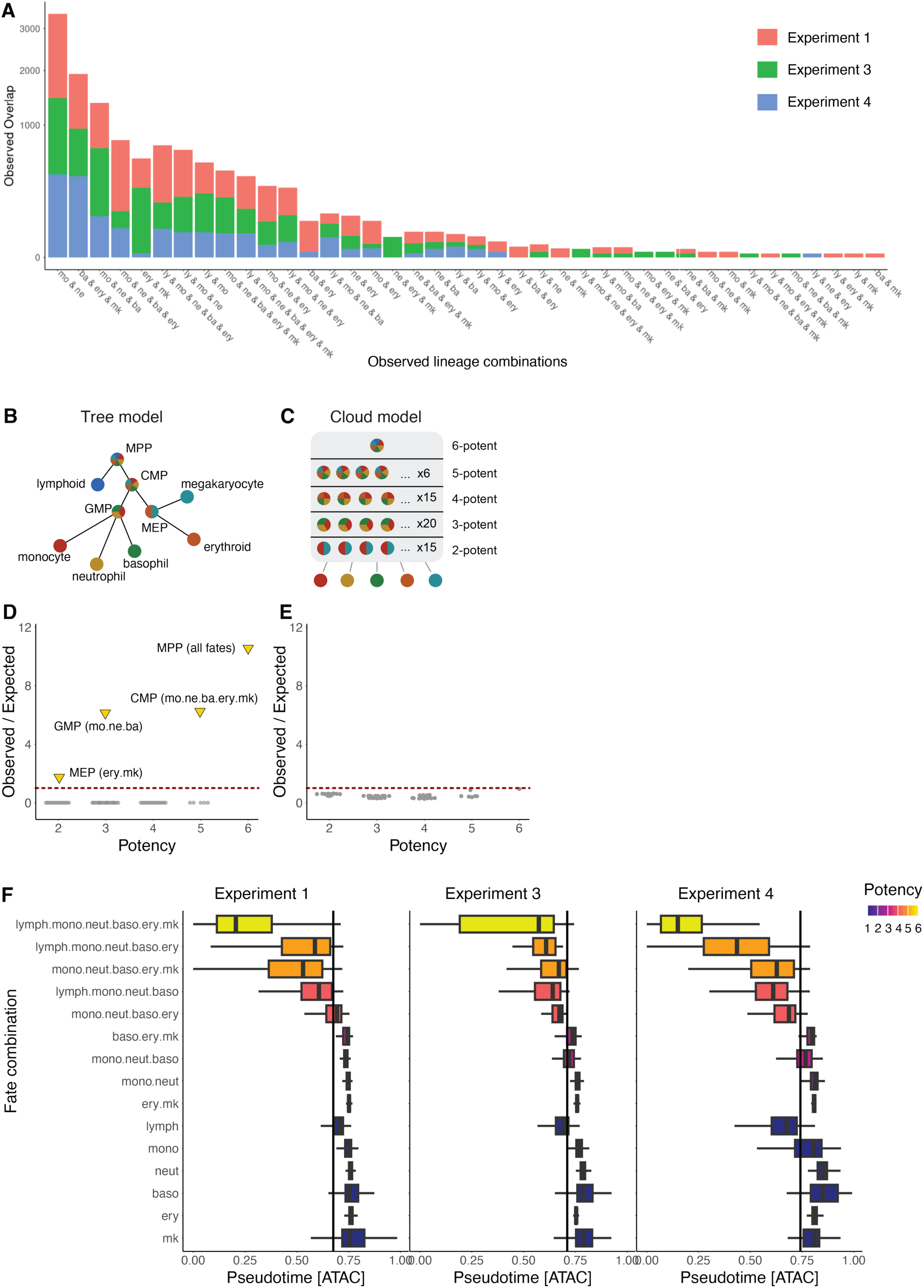
Emergence of lineage-restricted progenitors relative to proliferative activation. (A) Number of cells possessing the fate combination indicated on the x axis across replicates. (B,C) Schematic representation of the models used to generate the synthetic datasets. In the tree model (B), stem cells undergo discrete restriction steps during commitment, generating a defined set of oligo-potent states. The cloud model (C) lacks oligo-potent states, resulting in a large number of fate combinations observed. (D,E) Outcome of the statistical test on fate combination measured as ratio between observed and expected number of progenitors featuring a specific fate combination. The x axis enumerates the number of fates for the considered combinations. (D) In the tree dataset, the only significant combinations are the ones used to build the dataset. (E) The lack of significant combinations in the cloud model reflects the absence of oligo-potent progenitors in the synthetic dataset. (F) Distribution of pseudotemporal values for the enriched oligopotent states detected across replicates. The black vertical line marks the point in pseudotime at which half of the maximum fraction of cycling cells is reached.

**Figure S4.**
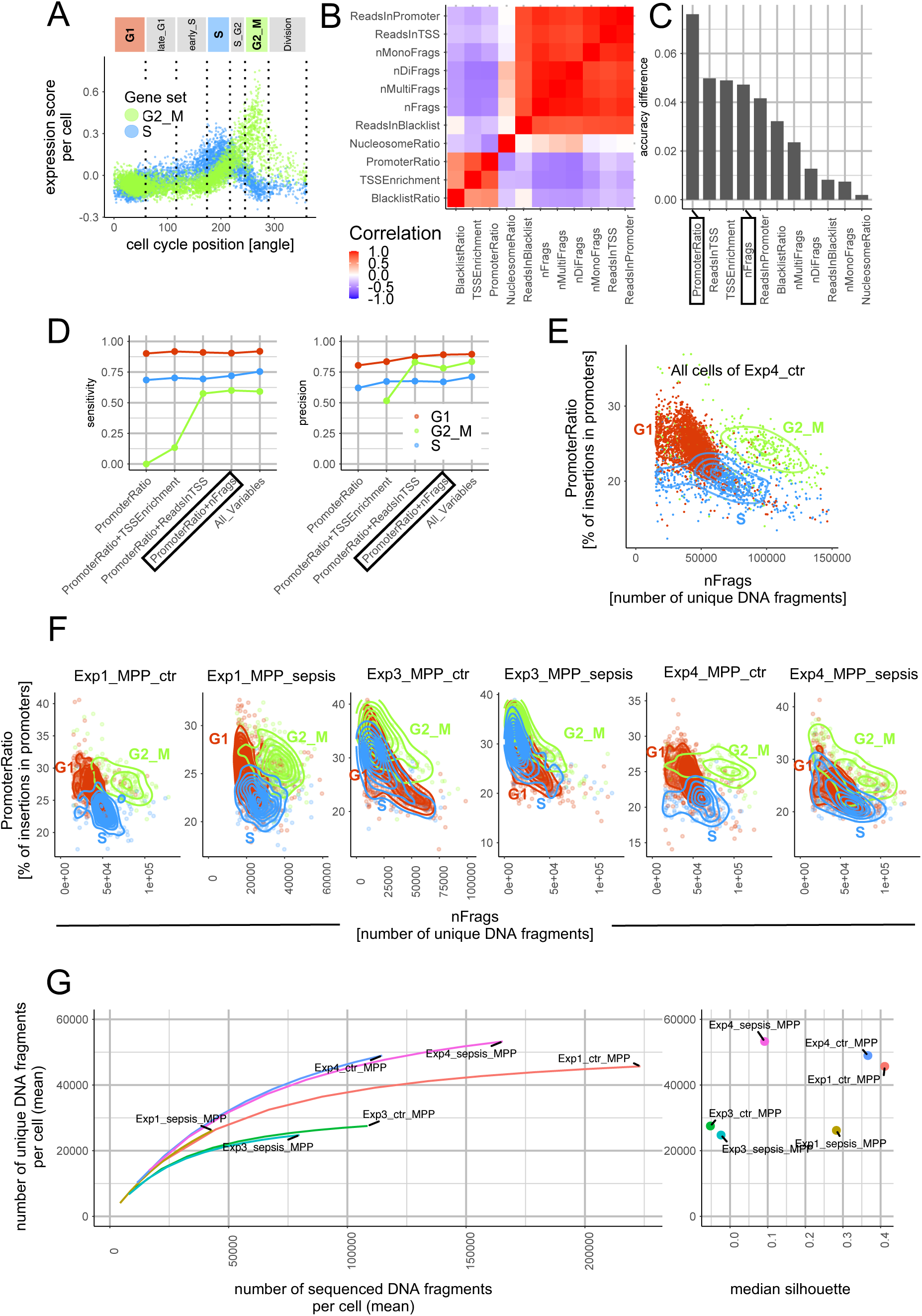
Pronounced cell-cycle signature in snATAC data from experiments with efficient DNA transposition. (A-E) Data of all hematopoietic progenitors from the untreated mouse of experiment 4. (A) RNA-based classification of cells into cell cycle phases. Cells are ordered along the x-axis according to cell cycle progression inferred by the *tricycle* software. For each cell, Seurat module scores (y-axis) are shown for two gene sets: One set consisting of genes that are specifically transcribed during S phase (blue) and another set with G2/M phase specific genes (green). (B) Heatmap shows Spearman correlations across all cells of experiment 4 (control) for cell-level chromatin accessibility meta-features computed by ArchR. These chromatin features were used to train a random forest model to predict the cell-cycle phase of a cell that was beforehand determined based on RNA data. (C) The importance of each feature for the accuracy of the prediction is indicated in the barplot which shows how the prediction accuracy drops if the trained classifier is applied to the training data upon permutation of the values of the respective feature across cells. (D) Dependence of random forest classifier performance in assigning cells to cell cycle phases on the selected sets of chromatin accessibility meta-features. Classifier performance is shown as sensitivity (left) and precision (right). (E) The total amount of accessible DNA (approximated by the number of unique fragments) and the percentage of accessible DNA mapped to promoter regions together separate nuclei by RNA-derived cell cycle phases. (F) Same as (E) but for MPPs of all mice. (G) Association of library complexity with the separation of cell-cycle phases by ATAC-features. Simulated sequencing saturation curves (Preseq software) for MPPs from distinct mice (shown in F). Mean silhouette scores per cell were computed using the RNA-based cell-cycle phase labels and cell distances based on ATAC-features (number of unique DNA fragments and fraction of transposase insertions in promoter regions).

**Figure S5.**
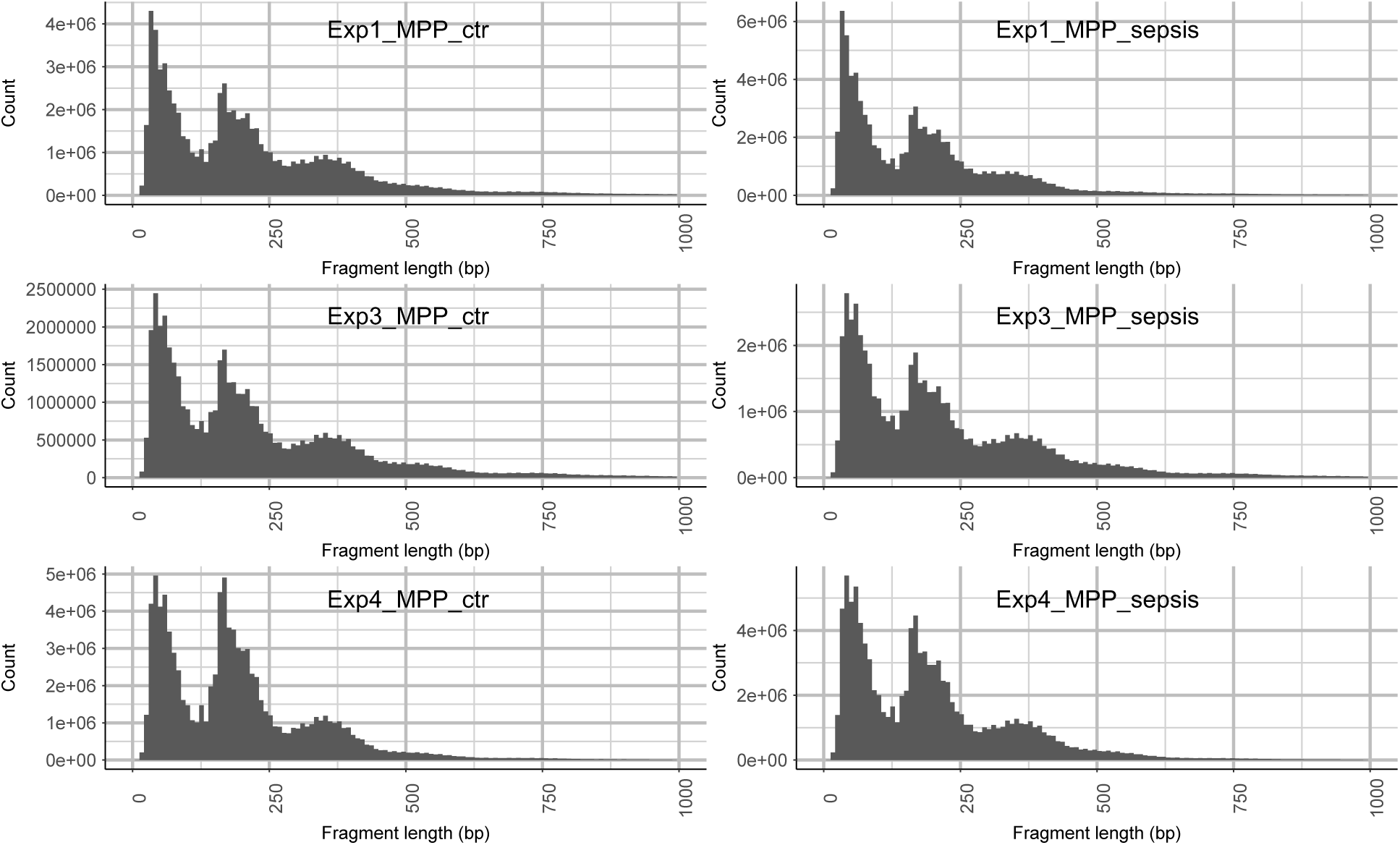
Tn5 transposition occurred in chromatin with intact nucleosome organization. Histogram of ATAC-seq fragment lengths for MPPs of all mice.

**Figure S6.**
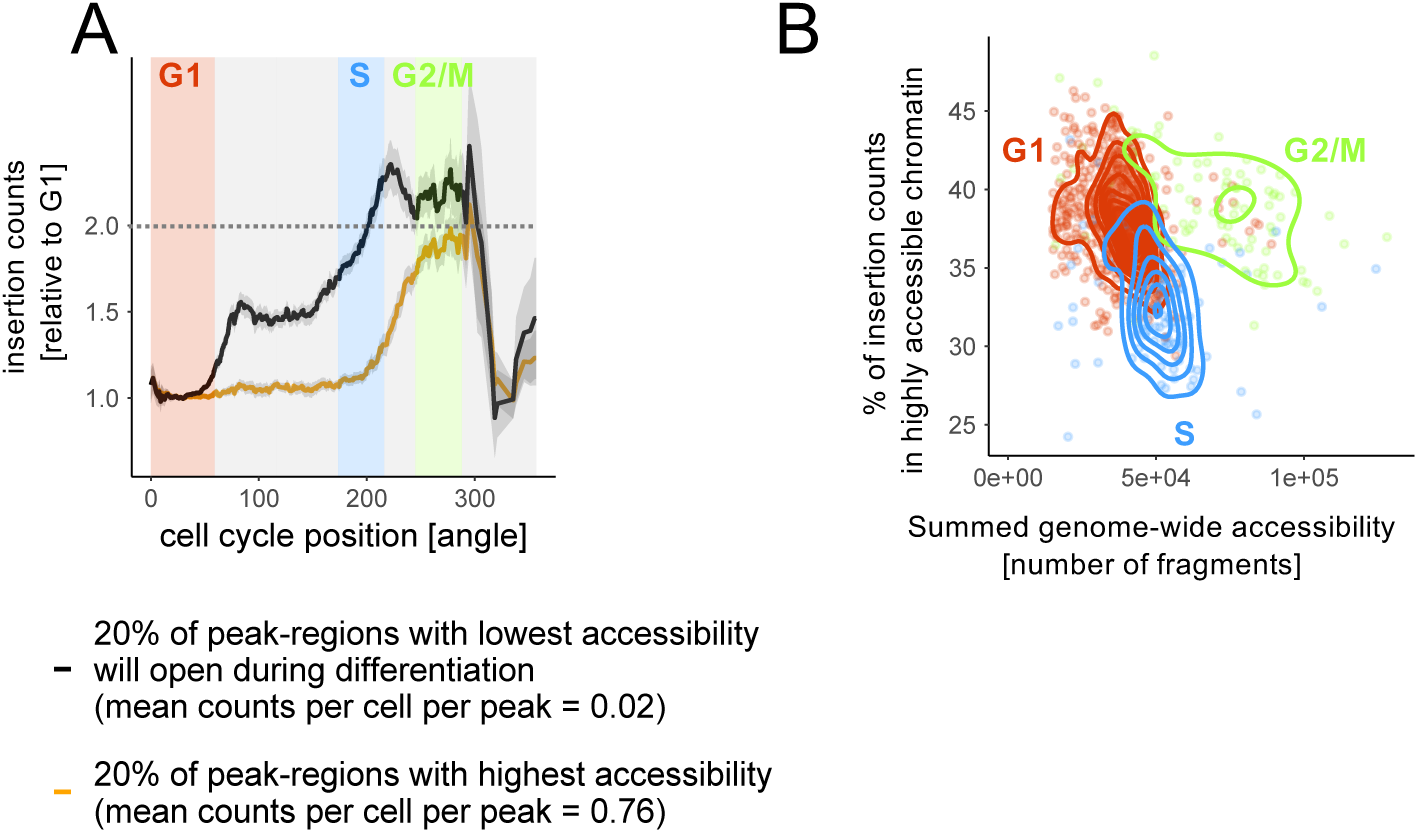
During S phase, uncommitted progenitors gain accessibility at lowly accessible loci (Experiment 1, control mouse). (A) Dynamics of chromatin accessibility (insertion counts) during cell cycle in transcriptionally uncommitted progenitors. Colors of curves encode the distinct baseline accessibilities at which the two types of genomic regions start in G1 phase. Y-values show the factor by which the initial accessibility has changed. Values represent moving averages (window size of 15°) of cells that were ordered by cell-cycle position based on their RNA profiles. (B) The total amount of accessible DNA per cell (x-axis) and the percentage of insertion counts in highly accessible chromatin (y-axis) together separate cells by their cycle phases that were determined based on their RNA profiles.

**Fig. S7.**
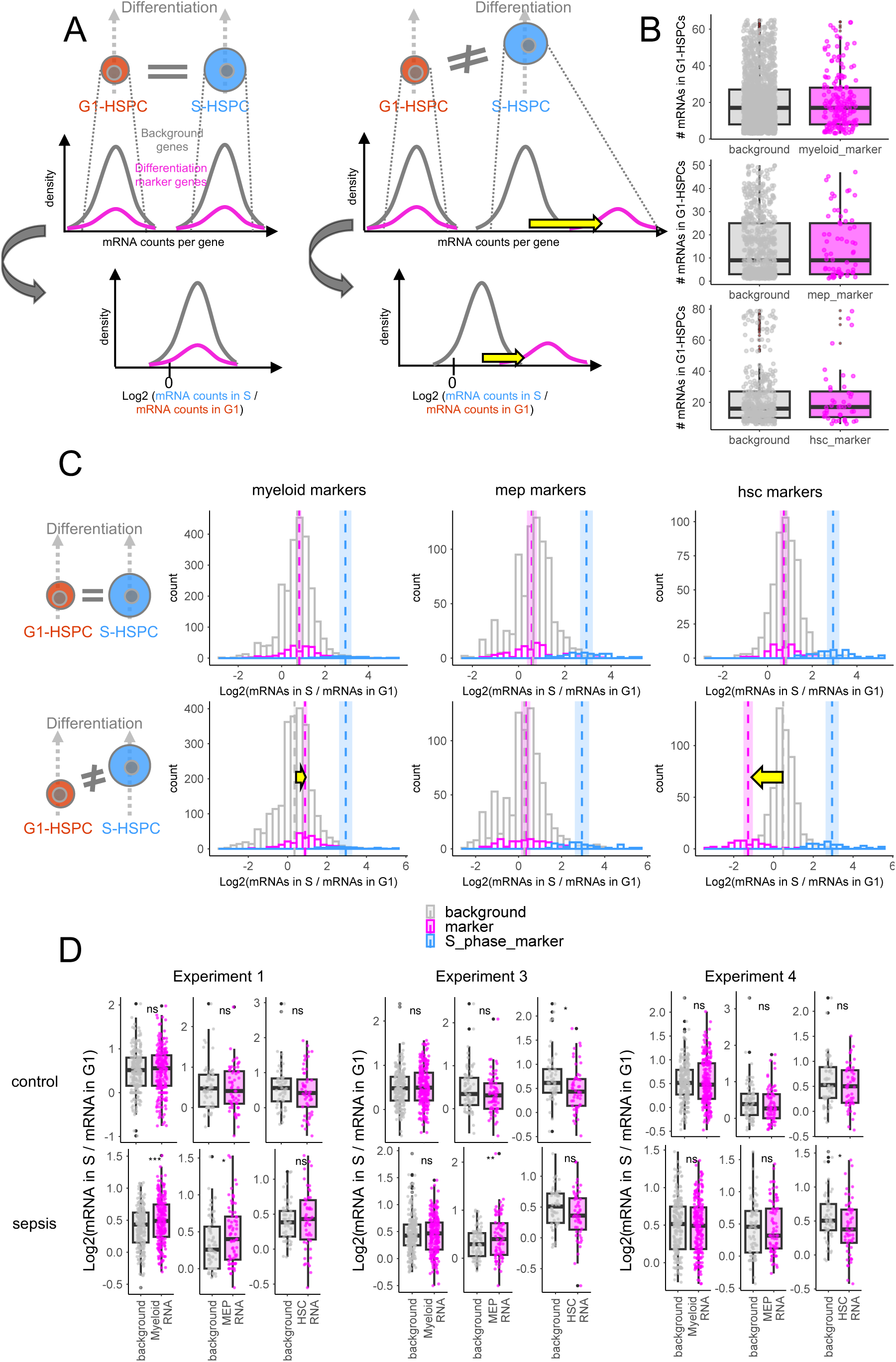
Validation: Uncommitted HSPCs in S phase are transcriptionally not more differentiated than G1-HSPCs to which they are compared. (A) Assumptions on differences in mRNA quantities between HSPCs of different cell-cycle phases. Comparison of equally differentiated HSPCs (left) and comparison of HSPCs in different differentiation states (right). (left, top) Distribution of marker mRNA* counts and of background mRNA** counts increase by the same amount while cells progress through cell-cycle. Therefore, fold changes of marker and background mRNAs between HSPCs of different phases are expected to be equal (left, bottom). If fold changes of marker mRNAs diverge from those of background mRNAs (right), HSPCs of compared cell-cycle phases are not in the same differentiation state. (B) Total counts of background and marker mRNAs in G1-HSPCs (Experiment 4, control). (C) (Top row) Comparison of background and marker mRNA fold changes between matched S-and G1-phase HSPCs that were compared in this manuscript (Experiment 4, control). Fold changes of S-phase marker mRNAs were added in blue as a reference for the size of fold changes. Vertical, dashed lines with shadows indicate means with 95% confidence intervals (bottom row) Negative example: Comparison between the most stem-like HSPCs and multipotent progenitors with highest expression of myeloid lineage markers. Cells were selected using Seurat gene module scores. (D) Test for differentiation state differences between matched G1 and S-phase HSPCs used for cell-cycle comparisons in this manuscript. Comparisons according to A-C for all samples. Dots represent genes. Marker genes are compared to background genes by a paired Wilcoxon test as one background gene was selected for each marker gene. Fold changes were computed based on raw mRNA counts. Comparable distributions of fold changes from marker and background genes suggest that G1 and S-phase HSPCs are equally differentiated / undifferentiated (indicated significance levels: p<0.0001: ****; p<0.001: ***; p<0.01: **; p<0.05*). *Marker mRNAs: mRNAs with changing abundancies during differentiation. ** Background mRNAs: mRNAs that are no markers but have same abundancies as marker mRNAs in G1 HSPCs.

**Figure S8.**
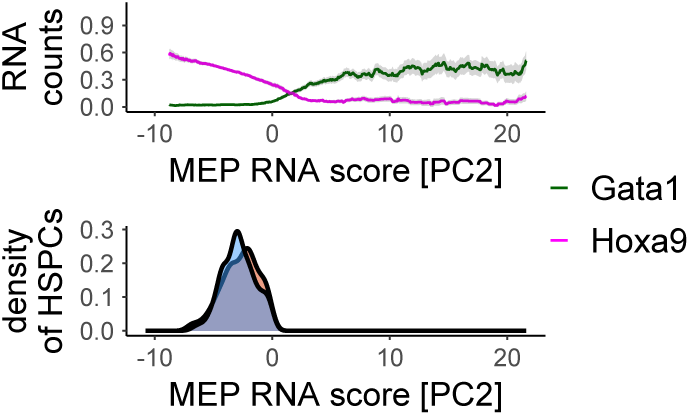
Increased transcription of Gata1 marks erythroid differentiation. Position of transcriptionally matched HSPCs in G1 and S phase along an RNA-based differentiation trajectory, ranging from transcriptionally uncommitted cells (PC2 < 0) to megakaryocyte/erythroid-committed cells (PC2 > 0). Colored lines represent moving averages of RNA counts. Grey ribbons indicate the standard error of the mean. Data shown are from the untreated mouse in experiment 4.

**Figure S9:**
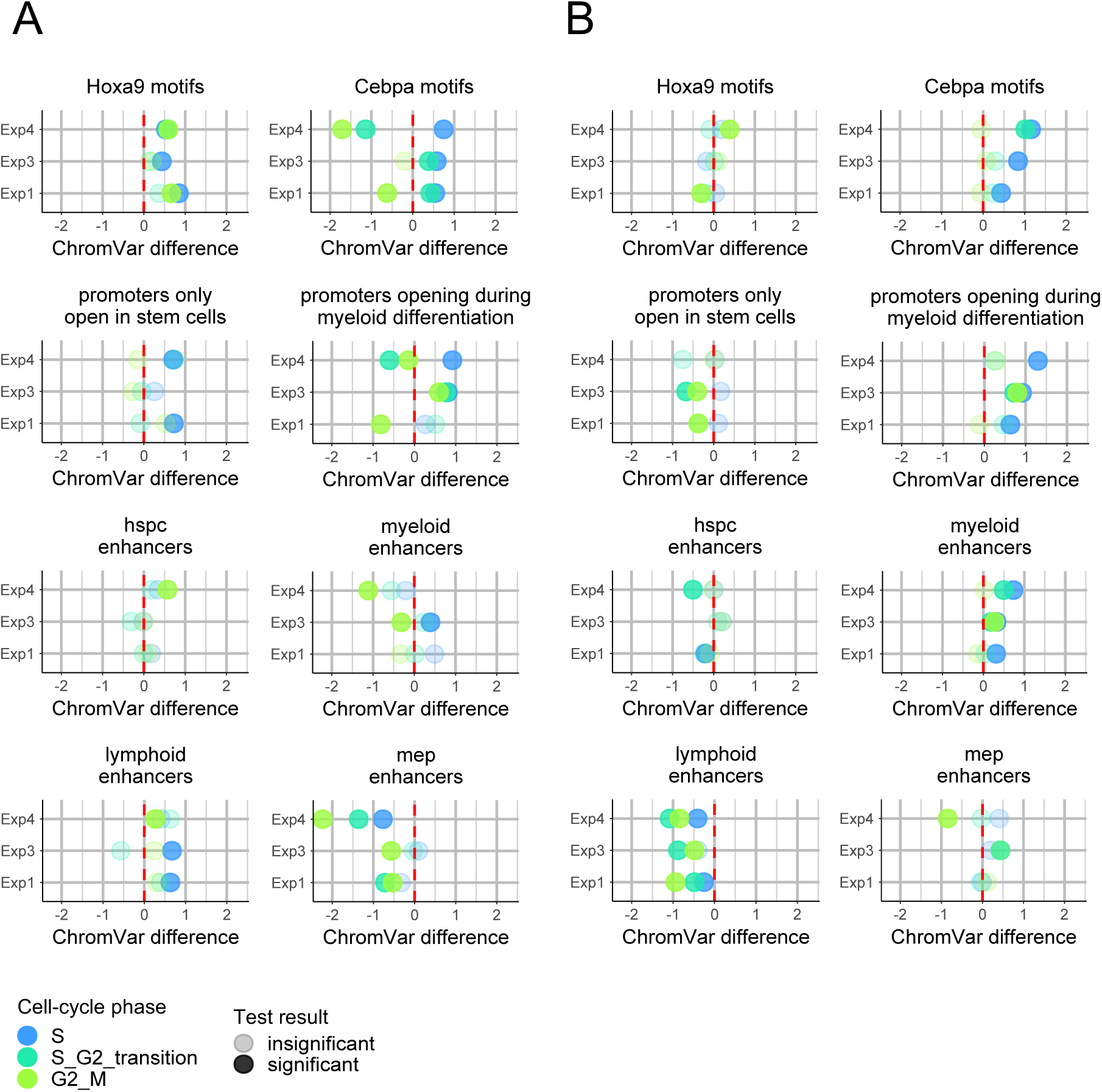
Accessibility changes during cell-cycle relative to G1-phase in transcriptionally uncommitted HSPCs. (A-B) Chromatin accessibility differences (x-axis) between HSPCs in G1 phase (vertical red dashed line) and HSPCs in other cell-cycle phases (indicated by color). Units on x-axis are differences in ChromVar z-scores. Values larger than zero indicate that HSPCs in the respective cell-cycle phase exhibit more accessible chromatin than HSPCs in G1. ChromVar z-scores of HSPCs progressing through cell-cycle were compared to scores of matched G1-HSPCs using a paired Wilcoxon test. Non-significant differences (p>=0.05) are more transparent. Cell-cycle associated accessibility changes are shown for untreated (A) and septic (B) mice across all experiments. HSPCs of the first experiment only consist of immunophenotypic MPPs whereas HSPCs of the other two experiments additionally contain low fractions of HSCs and short-term HSCs.

**Figure S10.**
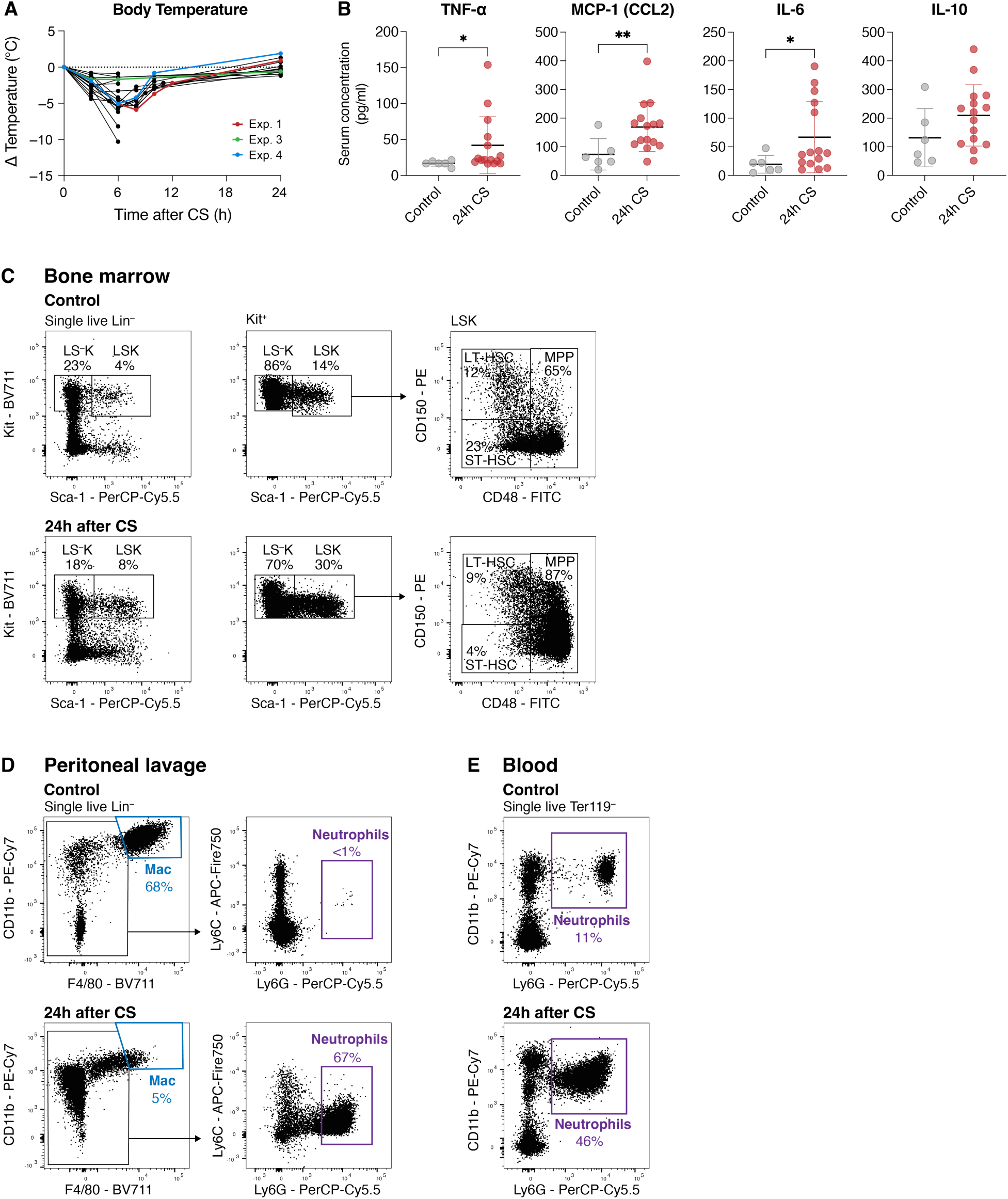
Cecal slurry treatment induces acute systemic inflammation with hallmarks of sepsis. (A) Changes in body temperature in individual mice after Cecal slurry (CS) treatment relative to temperature measured before CS treatment. Mice used for Multiome experiments highlighted in red (Experiment 1), green (Experiment 2), and blue (Experiment 4). (B) Serum cytokine levels (TNF-α, MCP-1, IL-6, IL-10) in untreated control mice and 24 hours after CS treatment. (C) Representative gating scheme used for sorting LS^-^K, LSK, LT-HSC, ST-HSC, and MPP in untreated control mice and 24 hours after CS treatment. (D) Representative flow cytometry gatings of changes in immune cell composition (reduction of peritoneal macrophages and increase of neutrophils) at the site of CS treatment analyzed by peritoneal lavage. (E) Representative flow cytometry gatings of neutrophil increase in the blood.

**Figure S11.**
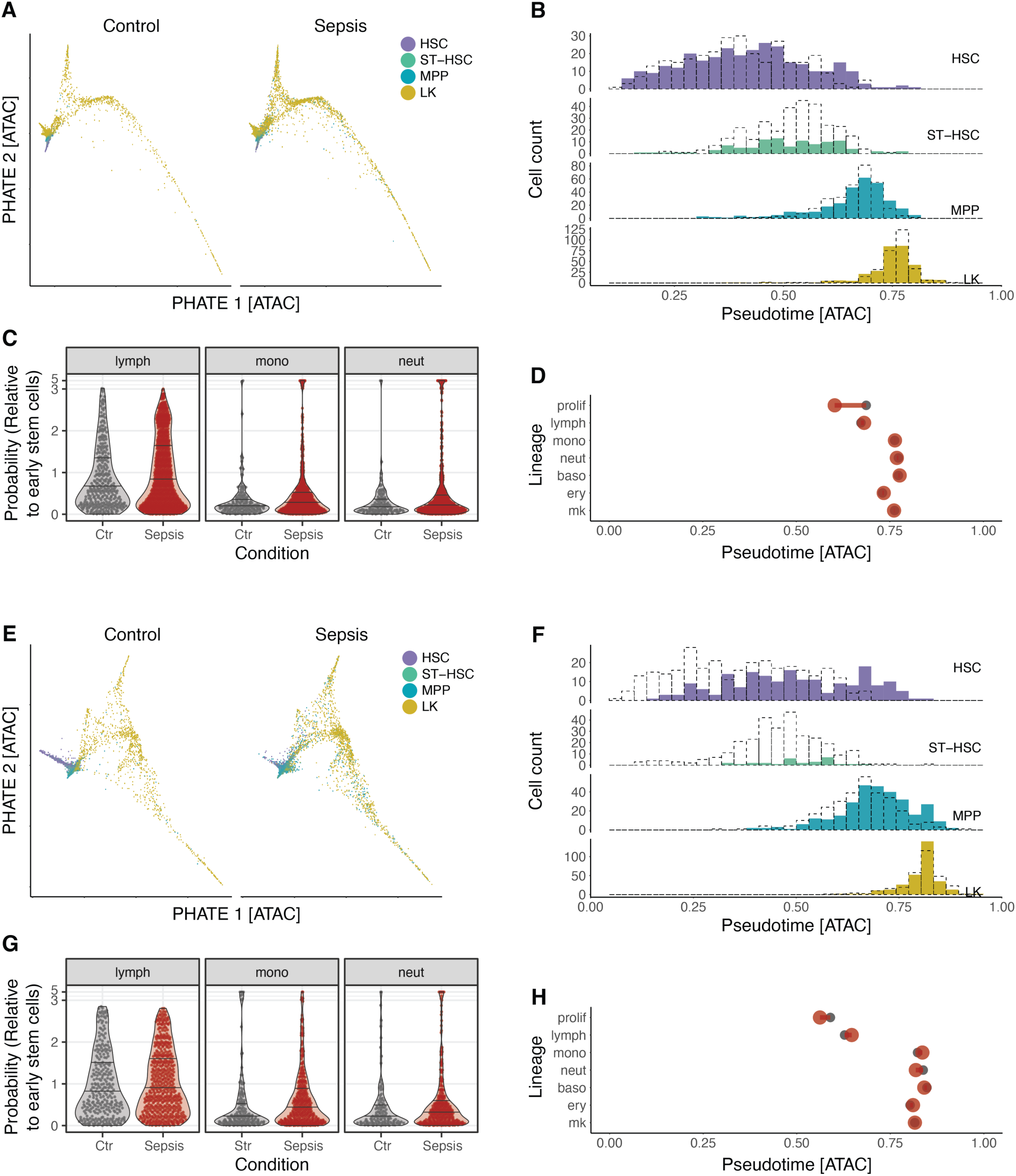
Emergency myelopoiesis during sepsis enhances proliferation and myeloid differentiation. (A-D) Analyses based on the control and sepsis mouse from experiment 3. (A) PHATE visualization of HSPCs under steady-state conditions (left) and in response to sepsis (right). Cells from the sepsis condition were projected onto the control dataset. Cells are colored according to FACS-sorted populations (HSC, ST-HSC, MPP, LK; see (B) for color code). (B) Density distribution of sorted cell populations (HSC, ST-HSC, MPP, LK) along pseudotime, comparing control (dotted lines) and sepsis conditions (filled histograms). (C) Distributions of lineage probability scores in LSK cells of control and sepsis conditions for the indicated lineages. Z-scores were computed by subtracting the mean probability observed in early stem cells and dividing by the corresponding standard deviation. Scores < 0 are not shown. (D) Pseudotemporal onset of cell cycle activation (top row) and lineage identity during sepsis. Onset was computed as half-maximum constant for the monotonic splines fitted to the pseudotemporal trend. (E-H) Same analyses for control and sepsis mouse from experiment 4.

**Figure S12.**
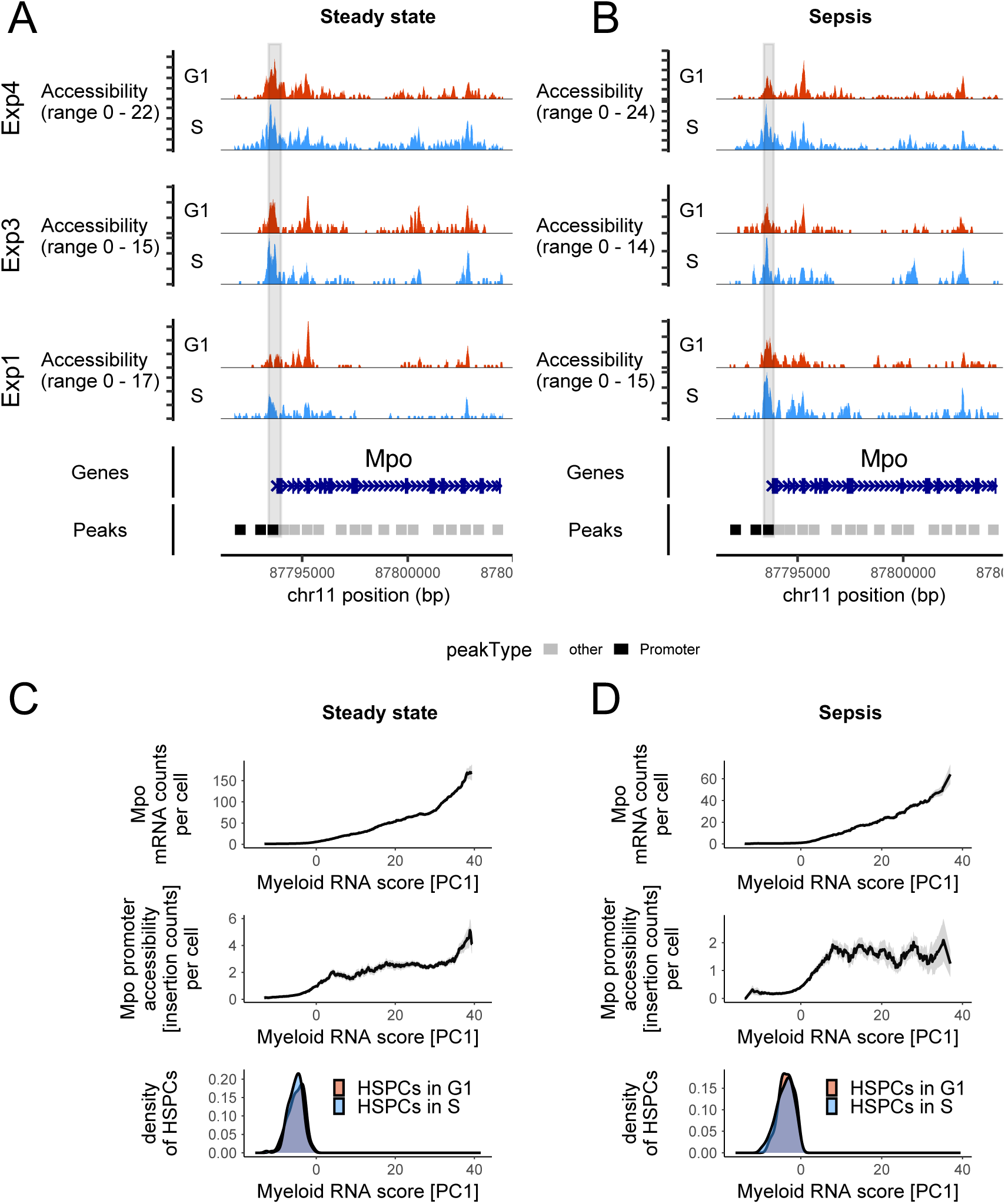
In uncommitted HSPCs, during sepsis, the promoter-region of *Mpo* becomes more accessible in S phase. (A,B) Centered moving sum of chromatin accessibility (Tn5 insertion counts) at the *Mpo* locus (moving window size = 100 bp). Tracks depict data from transcriptionally matched, uncommitted cells in G1 (red) and S (blue) phases for untreated (A) and septic (B) mice. Each marked peak spans 500 bp. Gray shading indicates the promoter peak overlapping the transcription start site (TSS). Tracks are unscaled, since moving sums were computed using equal numbers of cells per cell-cycle phase. (C,D) Expression of *Mpo* along the RNA-based differentiation trajectory, from transcriptionally uncommitted cells (PC1 < 0) to myeloid-committed cells (PC1 > 0). Promoter accessibility reflects insertion counts within the 500 bp promoter peak overlapping the TSS (shaded in A and B). All values are centered moving averages (window size = 4 units), based on the untreated (left) and septic (right) mouse from experiment 4. Gray ribbons denote the standard error of the mean. Cell-density curves mark the positions of uncommitted, transcriptionally matched G1 and S phase cells.

**Figure S13:**
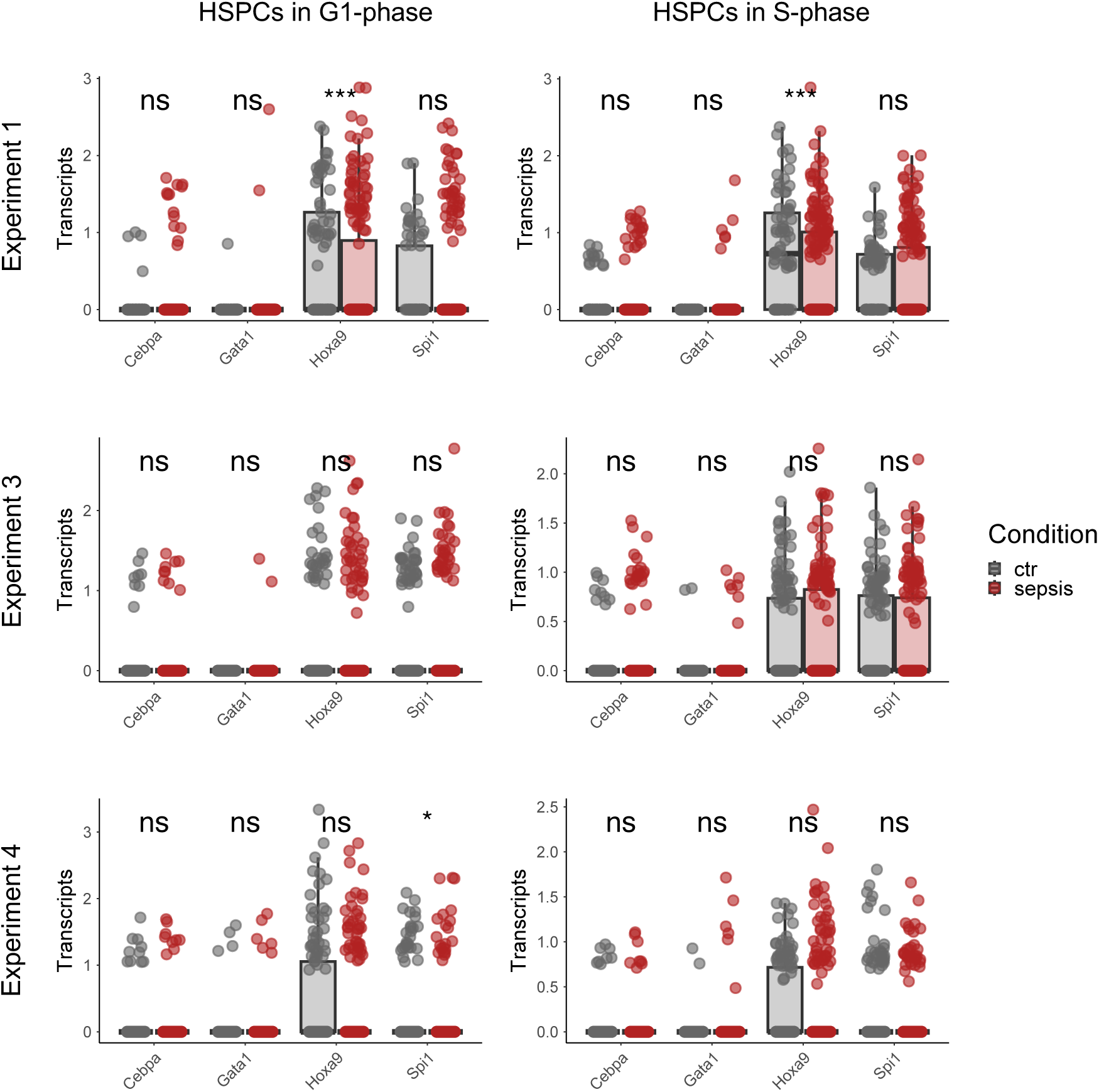
Major multipotency and lineage TFs do not exhibit clear expression changes in uncommitted HSPCs upon induction of sepsis. Normalized mRNA counts of TFs were compared between HSPCs of untreated and septic mice. Only transcriptionally uncommitted HSPCs that were used for comparing chromatin accessibility between G1 and S-phase (matched neighbors) were used. G1-HSPCs (left) and S-HSPCs (right) were separately compared between the control and septic mouse within each experiment. Stars indicate p-values computed by Wilcoxon test (p<0.001:***; p<0.05:*).

## RESOURCE AVAILABILITY

### Lead contact

Further information and requests for resources should be directed to and will be fulfilled by the lead contact, Thomas Höfer.

## Materials availability

This study did not generate new unique reagents.

## Data and code availability

The 10x Multiome data have been deposited in the Gene Expression Omnibus data base with the identifier GSEXXXXXX. These data are currently private and will be released upon publication in a peer-reviewed journal. Accordingly, the custom code used to analyze data will become publicly available on GitHub at the time of publication.

## ACKNOWLEDGEMENTS

We thank all members of the Höfer and Rodewald group for ongoing support and discussions, the Single Cell Open Lab for advice and support, Sven Schäfer for technical support, and the team of animal caretakers at DKFZ for support and expert animal husbandry. HRR and TH acknowledge core funding from the German Cancer Research Center (DKFZ) and the Sonderforschungsbereich SFB 873-B11 of the Deutsche Forschungsgemeinschaft.

## Author contributions

J.M., A.G., L.F. and T.H. conceptualized and designed the study. J.M., A.G., L.F., and N.C. planned the 10x Multiome experiments. L.F., performed sepsis experiments and cell sorting. J.M., and N.C. performed 10x Multiome experiments, N.C. prepared libraries for sequencing. J.M. and A.G. analyzed and interpreted the 10x Multiome data, L.F. analyzed flow cytometry and serum cytokine data. J.M., All authors interpreted the data and wrote the manuscript. T.H. and H.-R.R. supervised the study.

## METHODS

### Mice

Mice were kept in individually ventilated cages and under specific pathogen-free conditions in the internal animal facility at the DKFZ. No randomization or blinding was used for experiments. Only male wildtype C57BL/6J mice of age 12 to 16 weeks were used. All animal experiments were performed in accordance with institutional and governmental regulations and were approved by the authorities (Regierungspräsidium Karlsruhe, Germany).

### Sepsis induction and monitoring

C57BL/6J mice were euthanized, and the cecum was dissected to collect its content using sterile forceps and a spatula. The cecal content was then weighed and 100 mg were mixed with 0.5 mL sterile water. The mixture was sequentially filtered using sterile mesh strainers with pore sizes of 860 µm, 190 µm, and 70 µm. The final filtrate was mixed with an equal volume of PBS/30% glycerol, creating a final 15% glycerol stock of cecal slurry (CS), and aliquots were frozen at - 80°C. All experiments were performed using aliquots of the same CS stock. To avoid batch effects and storage stability, colony forming units (CFU) were determined regularly by spreading different dilutions of CS on agar plates prepared with brain-heart-infusion broth (BHI) that were incubated for 24 hours before counting colonies. CFU values remained robustly at 1.4 × 10^4^ CFU/µL (± 0.37 × 10^4^ SD) over three years. For sepsis induction, mice received 5 µL CS (i.e., 7.1 × 10^4^ CFU) per gram body weight as intraperitoneal (i.p.) injection. To minimize confounding effects related to sex or strong physiological differences, only male mice between 12 - 16 weeks of age were included.

Immediate effects of CS injection were observed by measuring the rectal body temperature 3, 6, and 24 hours after treatment. If body temperature decreased by ≥ 3°C, a body score was determined according to the appearance of the animal’s fur, eyes, and state of consciousness, and temperature was measured again after two hours. Animals were eliminated once the rectal body temperature decreased by ≥ 6°C or when the animal’s body score exceeded the threshold of ≥ 7. Serum cytokine levels were measured 24 hours after CS injection or at earlier end points. Bacteremia, systemic bacteria in the blood, was measured by collecting blood from the heart in EDTA-containing tubes. Whole blood was diluted by factors 2, 4, 8, 18, 32, 64, 128, and 256, and from each dilution a volume of 80 µL was spread on two agar plates prepared with brain-heart-infusion broth (BHI). Plates were incubated overnight at 37°C, colonies were counted after 24 hours, and the average colony number was calculated for each dilution with countable colony density from duplicates or triplicates to calculate the colony-forming units per microliter blood.

### Bead-based detection of serum cytokines

To measure serum cytokine levels, blood samples were collected from the facial vein into microtainer tubes containing a separation gel. After incubating for 30 min at room temperature, samples were centrifuged for 1.5 min at 8,000 × *g* to separate the serum fraction from solid blood cells. The sera were then transferred into polypropylene tubes and stored at -80°C. Prior to analysis, samples were thawed on ice and immediately processed using the bead-based cytokine detection assay LEGENDplex**^TM^**. Serum samples were analyzed in duplicates or triplicates (25 µL input per well) using 1:2 dilutions (controls and all time points beyond 6 hours post CS) or 1:10 dilutions (3 and 6 hours post CS) of sera. Initial incubation of sera with detection beads was performed overnight at room temperature by fixing the plate on top of a thermal shaker with constant shaking of 500 - 600 rpm. All remaining steps were conducted according to the manufacturer’s instructions with adjusted volumes to the reduced input volumes. Beads and the PE-based cytokine levels were measured by flow cytometry on an LSR Fortessa machine and analyzed with the LEGENDplex**^TM^**software Qognit.

### Fluorescent activated cell sorting

Bone marrow cells were collected from different bones of the skeleton (femora, tibiae, fibulae, pelvis, humeri, radii, ulnae and spine) by crushing bones in PBS/5% FCS using mortar and pestle. Cells were filtered through a 40 µm strainer, blocked for 20 min with purified whole IgG, and stained with lineage markers (CD4, CD8a, CD11b, CD19, Gr-1, NK1.1, and Ter119) for 30 min on ice. After washing (PBS/5% FCS), cells were incubated with magnetic Dynabeads for 1 hour at 4°C on a rotor. Cells in the supernatant that did not bind to magnetic Dynabeads were processed for further blocking and staining.

Before blocking and staining, single-cell suspensions were counted using ViaStain AOPI staining solution. The AOPI solution combines acridine orange, staining all nuclei (both live and dead cells) with green fluorescence, and propidium iodide, which stains nuclei of cells with compromised membranes (dead cells) with red fluorescence. Propidium iodide quenches green fluorescence emitted by acridine orange, thus, cells emitting only green fluorescence from acridine orange were identified as live cells and counted using the Cellometer Auto 2000 instrument. The entire available sample was further processed with blocking and staining. Cells were sorted with the LSR II Aria using a 100 µm nozzle and sorted cells were collected in 300 µL PBS/50% FCS collection buffer. LSK (Sca-1^+^ Kit^+^) or LT-HSC (Sca-1^+^ Kit^+^ CD150^+^ CD48^-^), ST-HSC (Sca-1^+^ Kit^+^ CD150^-^ CD48^-^), and MPP (Sca-1^+^ Kit^+^ CD150^-^ CD48^+^) were sorted from lineage-depleted bone marrow cells, as well as LK (Sca-1^-^ Kit^+^).

**Table 1.** Fluorescent antibodies used for flow cytometric activated cell sorting.

| Antigen | Clone | Fluorophore | Vendor | Dilution |
| --- | --- | --- | --- | --- |
| CD4 | GK1.5 | BV421 | BioLegend | 1:400 |
| CD8a | 53-6.7 | BV421 | BioLegend | 1:400 |
| CD11b | M1/70 | BV421 | BioLegend | 1:400 |
| CD11b | M1/70 | PE-Cy7 | eBioscience | 1:400 |
| CD19 | 6D5 | BV421 | BioLegend | 1:200 |
| CD48 | HM 48-1 | FITC | eBioscience | 1:400 |
| CD150 | TC15- | PE | BioLegend | 1:200 |
| F4/80 | BM8 | BV711 | BioLegend | 1:100 |
| Gr-1 | RB6-8C5 | BV421 | BioLegend | 1:400 |
| CD117 (Kit) | 2B8 | BV711 | BioLegend | 1:800 |
| Ly-6C | HK1.4 | APC/Fire™750 | BioLegend | 1:200 |
| Ly-6G | 1A8 | PerCP-Cy™5.5 | BD Pharmingen | 1:100 |
| NK1.1 | PK136 | BV421 | BioLegend | 1:200 |
| Sca-1 | D7 | PerCP-Cy5.5 | eBioscience | 1:200 |
| Ter119 | TER-119 | BV421 | BioLegend | 1:200 |

### Preparation of ATAC and RNA sequencing libraries from the same single nuclei

After being sorted into 50%PBS / 50%FCS solutions on ice, cells were immediately lysed to extract nuclei, using the low input lysis protocol (version) provided by 10X Genomics. Nuclei were inspected using a light microscope to ascertain that nuclei had a clean round shape indicating sufficient but not excessive lysis and to ascertain low levels of cellular debris in the suspension. Nuclei were counted upon Trypan Blue staining using an automated cell counter. If the obtained nuclei concentration strongly exceeded 10,000 nuclei per 5µl, the nuclei were further diluted to avoid excessive generation of doublets. Nuclei were immediately further processed as described in the Chromium Next GEM Single Cell Multiome ATAC + Gene Expression user guide. In the first experiment, HSCs, ST-HSCs, MPPs and LKs were separately loaded into distinct wells of the 10X chip J, whereas these populations were pooled for the following experiments to avoid batch effects arising from distinct ambient RNA backgrounds. After quality control by analysis of fragment length distributions, RNA and ATAC libraries were sequenced on Illumina NovaSeq6000. In each experiment we distributed libraries over a sufficient number of flow cells to roughly obtain 100,000 read pairs per cell for the ATAC-library and 50,000 read pairs per cell for the RNA-library. Libraries of samples processed in the same experiment were multiplexed and therefore sequenced on the same flow cells.

### Processing of raw sequencing data, computational cell filtering and normalization of RNA counts

FASTQ files were processed using CellRanger ARC version 2.0.0 with mouse reference genome version mm10. For each sample, the filtered gene count matrix and ATAC-fragments file produced by CellRanger were used to construct a Seurat^61^ object containing RNA and chromatin accessibility information for all cells. Specifically, we first created an ArchR^62^ object containing and referencing ATAC data for all barcodes putatively representing cells. The intersection of barcodes with ATAC information and of barcodes that were included in CellRanger’s filtered gene count matrix was subjected to quality control: Barcodes associated with low numbers of DNA-fragments (nFrags<2000) or with a low enrichment of fragment-ends mapped to transcription start sites (TSSEnrichment<6) or with low numbers of detected mRNA molecules per cell (nCount_RNA<1000) were excluded. These initial filter criteria retained most of the barcodes that were classified as cell-associated by CellRanger, thereby guaranteeing that retained barcodes represented cells while avoiding extensive cell-barcode exclusion that might hamper doublet detection. Putative doublets were identified based on log-normalized RNA counts using scDblFinder^63^ in the cluster-dependent and random mode. Barcodes predicted to represent doublets by both approaches were excluded. Upon removal of doublets, barcodes with fewer than 15,000 detected DNA-fragments were excluded to ascertain high resolution of chromatin accessibility. This threshold was adjusted to 7,000 for samples of the third experiment in order to retain a comparable fraction of barcodes. From here onwards we refer to the remaining barcodes as cells. The RNA count matrix as well as a meta-data table containing the cells’ values of ATAC quality measures computed by ArchR were stored in a Seurat object. Next, RNA data were normalized for downstream analyses: For each gene, the relationship between its number of mRNAs per cell and the cell-specific RNA library size was modeled using SCTransform^64^. The method returns the deviations of a cell’s observed mRNA counts from their expected values, also referred to as Pearson residuals.

### RNA-based assignment of cells to cell-cycle phases

Cells were assigned to cell-cycle phases using a two-step approach based on RNA data. First, they were arranged in a circular progression through the cell-cycle with the Tricycle software^65^. Then, Seurat’s cell-cycle scoring method was used to assign phases to different sections of the circle. Specifically, we used Tricycle’s pre-built loading matrix, which contains values indicating each gene’s contribution to two cell-cycle dimensions. To generate a two-dimensional circular representation of the cells, we multiplied their SCTransform-derived log-normalized RNA counts by this loading matrix. Each cell’s position in the cell cycle was then determined by its angle relative to the center of the circle. To identify sections within this circular cell-embedding which correspond to specific cell-cycle phases, we plotted Seurat’s RNA module scores of marker genes for S phase and G2/M-phase against cell-cycle angles. Gene sets with S and G2/M phase markers were loaded from the Seurat package (sets were originally derived from Tirosh et al.^66^). S phase and G2/M phase marker scores exhibited clear peaks and threshold-angles marking S and G2/M-phases were set tightly around these peaks to guarantee high precision of cell-cycle phase assignment. A section with high purity of G1-cells was inversely identified by simultaneously low S phase and G2/M-phase marker scores. To keep the fraction of false positive cells within each cell-cycle phase low, the described approach left intermediate sections in the circular cell-cycle embedding separating G1, S and G2/M-phase sections. As the embedding clearly resolved a progressive increase of S phase marker transcripts from the G1 to the S phase section, we further split this section into a late G1 and early S phase section. Sections were finally annotated as shown in Figure S4A. Inspection of transcript counts of individual markers genes along Tricycle-derived cell-cycle angles further supported the use of selected thresholds for accurate assignment of cells to cell-cycle phases. The same thresholds were applied to all samples. For Quantification of cell-cycle phase fractions along pseudo-time, each cell was assigned to one major phase (G1, S or G2_M). In this case, threshold-angles were selected to align best with Seurat’s CellCycleScoring based phase prediction (G1 <= 116° < S <= 231°< G2_M < 345° <= G1).

### Construction of RNA-based cell embedding using lineage markers

To perform dimensionality reduction, we selected genes whose transcript abundances changed during differentiation independently of the cell-cycle state. This approach ensured that the lower-dimensional embedding primarily reflected the differentiation trajectory rather than cell-cycle effects. By avoiding regression of cell-cycle effects, we prevented the potential reduction of Pearson residuals of differentiation markers. This was important because, in hematopoiesis, differentiation is accompanied by increased proliferation, leading to correlations between the expression of cell-cycle genes and lineage marker genes. For selection of lineage marker genes, we first identified genes with highly variable transcript counts based on the variance of their Pearson residuals, as computed by SCTransform. We then used the Pearson residuals of these genes to apply PCA and PHATE^30^ and generate coarse-grained cell clusters, which clearly distinguished cells based on their differentiation state. To extract markers of these differentiation clusters, we applied Seurat’s FindAllMarkers function. To further reduce the risk of retaining cell-cycle markers in this selection process we performed marker selection only on cells in G1 phase (cell-cycle annotation see below). The procedure yielded approximately 700 genes that were strongly enriched for differentiation markers while being largely free of known cell-cycle-related genes (S and G2/M phase related genes were loaded from the Seurat package; sets were originally derived from Tirosh et al.^66^). Gene selection was performed using data from the untreated mouse in Experiment 4. To ensure consistency across analyses, the same set of selected genes was used for dimensionality reduction in all other datasets. This set of differentiation marker genes was then used to generate the final cell embedding via PCA and PHATE. Only data of experiment 1 required adjusted processing steps for dimensionality reduction because of batch effects between LTs, STs, MPPs and LKs that were processed separately and hence exhibited distinct ambient RNA profiles. Instead of applying PCA and PHATE, as done for all other experiments, cells of experiment 1 were mapped into the PHATE space of experiment 4 using Seurat’s FindTransferAnchors and TransferData functions (control 1 to control 4; sepsis 1 to sepsis 4).

### Cell type annotation

In the first experiment, the FACS-based identity of hematopoietic progenitor subsets (LTs, STs, MPPs, and LKs) of one normal mouse and one septic mouse were retained throughout the single-cell sequencing process. This was not the case for the remaining experiments because all cells that came from the same mouse were loaded into the same well of the 10X Genomics Next GEM chip to avoid batch effects caused by cell type-specific ambient RNA profiles. Therefore, cell type annotation had to be done computationally for samples processed with this setup. To this end, we computationally aligned cells of samples with known immune-phenotype (experiment 1) and cells without labels using Seurat’s FindTransferAnchors function. This function projected data of experiment 1 into the PCA-space of the other experiments. Upon alignment, we then transferred labels from experiment 1 to cells of the label-free samples using Seurat’s TransferData.

### Construction of ATAC count matrices

We constructed distinct types of ATAC count matrices, including one that contained all genomic regions with significant accessibility and two subset matrices filtered for promoter or enhancer regions. Counts in ATAC-matrices describe the amount of enzymatic cuts/insertions that are detected within a particular genomic region in a cell.

To select all genomic tiles (each 500 base pairs long) with significant accessibility in at least a subset of cells, we used the addReproduciblePeakSet function of ArchR. All tiles returned by this function were used to construct the full ATAC-matrix, called peak matrix. Through the addReproduciblePeakSet function, peak calling was conducted using the MACS2^67^ software (version 2.2.9.1). Each obtained peak region had a length of 500 base pairs (bps). Peak calling was conducted based on data of the untreated mouse from experiment 4, as these data exhibited the highest ATAC-signal coverage per cell. For consistency, the same peak-regions were then used to construct ATAC-count-matrices for all other samples/mice.

ArchR’s peak annotations—based on the mm10 reference genome that was loaded via the Bioconductor R-package BSgenome.Mmusculus.UCSC.mm10 (version 1.4.3)—were used to extract the subset of peaks that overlapped with promoters. ArchR defines promoters as regions ranging from 2000 bps upstream to 100 bps downstream of annotated transcription start sites. To select peaks located in putatively regulatory enhancers, we extracted the subset of called peaks that overlapped with hematopoietic enhancers reported by Lara-Astiaso et al.^28^.

### Computing genome-wide accessibility at transcription factor binding motifs

Transcription factor binding motif accessibility was computed at single-cell resolution using ChromVar^47^. To this end, we first downloaded information on mouse transcription factor binding motifs from the 2024 release of the JASPAR^68^ database. Downloaded information was used to annotate ATAC peak regions by presence of binding motifs in their DNA-sequences using Signac^69^’s AddMotifs function. Finally, Signac’s RunChromVAR function was executed to compute a TF by cell matrix with ChromVar z-scores. Such a score expresses a cell’s accessibility in peak regions with binding motifs of a particular TF, relative to the background accessibility-signal in that particular cell.

### Construction of ATAC-based cell embedding

To generate low-dimensional ATAC-embeddings with PHATE, we first applied standard preprocessing to the chromatin enhancer peak matrix. We normalized counts using term frequency–inverse document frequency transformation (Signac::RunTFIDF) and identified informative features (Signac::FindTopFeatures). Dimensionality reduction was performed with singular value decomposition (Signac::RunSVD), retaining components whose correlation with sequencing depth (log10 number of fragments) was below 0.75. The selected components were then used as input for PHATE (phateR, version corresponding to Python PHATE), which was run with 30 nearest neighbors, embedding dimension set to 2, and a fixed random seed to ensure reproducibility.

### Trajectory inference on ATAC features

#### Subsampling

To ensure high number of LSK cells, the FACS experiment was designed to enrich for LSK populations prior to sequencing. As a result, the sequencing dataset contains disproportionately more primitive cells with respect to physiological proportions in the bone marrow. This sampling bias can affect trajectory inference by creating artificially dense regions in the neighborhood graph. To avoid this, trajectory inference was performed on a subsampled dataset that maintained the relative ratios between populations observed in the FACS experiment.

Additionally, we noticed a few cells with high pseudotime that did not express lineage markers and were poorly connected to the main partition in the data. To remove them, we computed diffusion pseudotime (ATAC features) and lineage scores (RNA features) on the full dataset. Cells with DPT values exceeding those of terminal lineage cells but lacking high lineage marker expression were classified as outliers and removed from downstream analysis.

#### Identification of tip stem cells

To establish the root of differentiation trajectories in the subsampled dataset, we first performed singular value decomposition (SVD) on the TF-IDF-normalized enhancer accessibility matrix, filtering out components that were highly correlated (|r| > 0.75) with sequencing depth (log10-transformed fragment counts).

LSI embeddings were fed into the DiffusionMap algorithm^29^. To identify the stem cell population, we calculated Pearson correlation coefficients between each diffusion component and transcriptional stemness scores. The cell with the highest projection on the component maximally correlated with stem score was used as root of diffusion pseudotime (DPT) computation on the subsampled

#### Detection of macrostates

We used CellRank^70^ to identify discrete macrostates to use as “endpoints” during the next steps. Using the previously computed diffusion pseudotime, we instantiated a PseudotimeKernel object(threshold_scheme=“soft”) to model transition probabilities between cellular states. We used the GPCCA decomposition method to coarse-grain the cell-cell transition matrix onto the macro-state level.

To identify macrostates representing terminal lineage fates, we computed median lineage scores for each macrostate, and retained as terminal states the ones whose median score exceeded a defined threshold (standardized score > 1.75) for any lineage.

Finally, we used compute_fate_probabilities to computed absorption

#### Projection of results to full dataset

We projected the fate probabilities and diffusion pseudotime values from the subsampled dataset to the full dataset using support vector machine (SVM) regression. LSI components of cells of the subsampled dataset cells were used as training data, with pseudotime and probabilities as target variable. A SVM implementation from the package e1071 was trained with fixed parameters (γ = 0.1, cost = 10), achieving extremely high precision on training data (R2 > 0.99 for each task). The trained SVM models were then used to predict pseudotime and probabilities to the full dataset, with the additional costraint of clipping values to the [0,1] range.

### Monotonic spline fitting using ATAC-based pseudotime

Monotonic curves were fit using the penalized R package with L2 regularization. We used I-splines as the basis functions, generated via the iSpline() function from the splines2 package^71^, which ensures monotonicity when coefficients are constrained to be positive. The fitting procedure utilized optL2(positive = T, fold =10) to enforce positive coefficients and used cross-validation to select the best regularization coefficient. The half-maximum point (t_half) was calculated from the fitted curve to identify the pseudotemporal position where the curve reaches 50% of its maximum. To compute proliferation trends, cells were binned into 40 pseudotime bins containing a similar number of cells (function cut_number). For each bin, we calculated the proportion of cycling cells (i.e., cells not assigned to G1) in the tricycle assignment.

Differently from the proliferation trend, lineage probability trends are characterized by the co-occurrence of upwards and downwards trends, prompting us to use a two-step approach to isolate and parametrize only the upwards trend:

For each lineage, probability trends along pseudotime were fit using a two-step approach to better parameterize the curves: First, we fitted a monotonic spline to cells with probability values above the median probability among early progenitor cells. In a second refinement step, we selected points that were either close to the fitted curve (residuals > -0.025) or showing very high probability (p>0.5), and fit a second monotonic spline to these points. In this way, we made sure to only use cells in the upper side of the bifurcating plot, thus capturing the earliest events related to lineage emergence.

### Detection of lineage bifurcations using fate potential model

To assign lineage potential to each cell, we implemented a two-step procedure based on the bifurcating shapes observed in pseudotime-probability scatterplots. Initially, cells are labelled as lineage-potent for a lineage if their probability exceeds the median probability among early progenitor cells (i.e. cells with pseudotime lower than a threshold). The outcome of this step is a combination of fate potentials for each cell. Most early pseudotime cells are assigned all fates, while for higher pseudotemporal values the number of available fates gradually lowers to one.

In a successive refinement step, we reasoned that successive bifurcation in the pseudotime-probability trends would render necessary an adjustment of the probability threshold used to define fate. To understand why, it’s useful to illustrate the example in Figure 2B. In this example tree, the lineage A branches off before fates B and C. Due to the normalization of probabilities and due to the loss of potential for fate A, the probability for fates B and C after the branch A has split off becomes higher. If the probability threshold is not adjusted to account for this branching event, cells will be artificially assigned a bi-potent BC state even after B and C split.

To avoid artifacts in successive branching events, each combination of fate potentials was iteratively considered as root for downstream populations, progressing from high to low potency states (6→5→4→3 potential lineages). For each previously defined state containing n fates, we identified downstream cells with n-1 fates and defined an updated threshold computed as the mean probability of the relative root, thus removing the lineage label to cells with probability lower than the updated threshold.

### Statistical testing of fate combinations

To discern fate combinations that could correspond to biologically relevant states, we used SuperExactTest^72^, an R package that performs statistical testing of multi-set intersections. In this way, detected fate combinations are compared against a null model in which fate labels are assigned at random.

To evaluate this method, we generated two synthetic models with controlled properties:

- The tree dataset was constructed by directly specifying cell states corresponding to a classic hierarchical differentiation paradigm,
- The cloud dataset was generated by randomly shuffling lineage assignments in the tree dataset.

When tested on the “cloud” synthetic dataset (see main text), the test does not detect any fate combination to be statistically significant, as expected by random shuffling.

Importantly, multiple combinations were detected as significant in the tree dataset, far exceeding the number of progenitors states we input. By investigating the enriched combinations, we concluded that a progenitor state, defined by a fate combination (e.g.: lymph.mono.neut.baso.ery.mk) will cause any combination that is a subset of the original one to be enriched too, despite the lack of a defined progenitor state.

To avoid overestimation of progenitor states, we devised an iterative version of the multiSet intersection test: for each number of fates (6→5→4→3), higher-order progenitor cells are removed prior to testing. Applying this procedure to the tree data recovered the progenitor states used to construct the dataset.

The iterative version of the multiSet intersection test was performed on the three control replicates. Significant combinations were defined based on effect size (mean FE > 1.5) and significance (p.value < 0.05 in at least one replicate).

### Projection of sepsis using ATAC features

#### Merge conditions

To enable direct comparison between control and sepsis samples, we concatenated and re-performed normalization in both expression and chromatin assays.

### Project trajectory inference results from control

To evaluate sepsis-induced changes in differentiation trajectories, we projected trajectory inference results from control samples onto the merged dataset containing both control and sepsis cells. We used Seurat’s FindTransferAnchors function with the control cells as reference. Using TransferData, we mapped diffusion pseudotime values and lineage-specific fate probabilities from the control-only analysis to all cells in the merged dataset. The same transfer approach was applied to dimensionality reduction coordinates (UMAP, PHATE) to enable visualization of all cells in a consistent coordinate system.

### Identification of cell-level ATAC meta-feature associated with cell-cycle

Genome-wide cell-cycle associated changes in ATAC signals were detected by testing which genome-wide ATAC features, computed by ArchR, contained information on the cell-cycle state that was inferred from RNA data as described before. All considered ATAC features are shown in (Figure S4B). Cells were split into a training (70%) and a test set (30%). A random forest classifier (R-package Ranger^73^; version 0.14.1) was trained to predict cell-cycle phases based on ATAC-features. To optimize the performance of the random forest model, a hyperparameter scan was performed and the parameter values that yielded the highest classification accuracy for the test data were used for model training. To identify the ATAC features that were most and sufficiently informative for the classification, we inspected the feature-importance scores returned after model training (Figure S4C). The importance score of an ATAC-feature quantifies how much this feature contributes to the classification accuracy of the trained model. As the *importance* parameter of the *ranger* function was set to *permutation*, the contribution of each ATAC-feature to the accuracy of the trained model was estimated by permuting the feature values across cells and recording the observed difference of classification accuracy. Computed importance most reliably reflects a features information on the cell-cycle phase if features are not strongly correlated. As our set of ATAC features contained two clusters of highly correlated features (Figure S4B), we primarily selected features based on importance, however, if importance scores were similar across correlated features, we chose the feature that was most representative for its cluster (most correlated with the other features of the same cluster). We trained the random forest model on combinations of prioritized features and found that the maximum classification performance, expressed by sensitivity and precision (Figure S4D), is achieved by using *PromoterRatio* in combination with *nFrags* or another feature that is strongly correlated to nFrags and thus genome-wide accessibility (like *ReadsInTSS)*.

### Comparison of chromatin accessibility across HSPCs in different cell-cycle phases

#### Definition of transcriptionally uncommitted HSPCs

From LTs, STs or MPPs, we removed these cells that contained elevated amounts of lineage-specific transcripts. All retained cells were considered to be transcriptionally uncommitted. To this end, we applied the same PCA rotation matrix (derived from experiment 4) to RNA Pearson residuals of all samples. The obtained principal components (PCs) 1 and 2 fully aligned with myeloid and megakaryocytic/erythroid differentiation respectively. In this two-dimensional representation LTs, STs and MPPs were mainly located in a region defined by PC1 and PC2 coordinates larger than zero (note that PC-values in Figure 5B, 6F, S8 and S12 were multiplied by -1). Cells outside of this region in PCA-space were re-labeled as committed cells.

#### Selection of promoter peaks with differentiation-dependent accessibility

To identify promoter peak regions with distinct chromatin accessibility between HSCs and cell states of early lineage commitment, we conducted differential testing using Seurat’s FindMarkers function. Markers were selected based on data of the control mouse from experiment 4 because this sample exhibited the highest resolution of chromatin accessibility per cell. Only cells in G1 phase were used to reduce the risk of confounded results. To identify promoter peaks that open immediately during commitment to the myeloid or MEP lineage, we respectively compared uncommitted cells (RNA-PC1 > 0; RNA-PC2 > 0) with cells slightly committed to the myeloid (−10 < RNA-PC1 < 0; RNA-PC2 > 0) or MEP lineage (RNA-PC1 > 0; -10 < RNA-PC2 < 0) (note that PC-values in Figure 5B, 6F, S8 and S12 were multiplied by -1). Only those markers that maintained increased accessibility until the most mature states were retained. RNA-based PCs were used for selection of early lineage committed states because the first two PCs clearly ordered cells by myeloid and MEP differentiation as shown in Figure 5B and Figure S8. As PCs are linear combinations of transcript counts, selection of cells by PC-thresholds corresponds to the assignment of cells to differentiation states by the weighted sum of differentiation markers. As uncommitted cells also contained LMPPs, identified myeloid markers were not contaminated by lymphoid markers. To detect promoter peaks that are specifically accessible in HSCs and close during lineage onset, we conducted the inverse comparison but this time only used tip stem cells instead of all uncommitted cells. This was necessary to avoid calling of lymphoid markers due to the presence of LMPPs. To this end, we increased the thresholds of PC1 and PC2, and removed LMPPs based on a threshold for PC4. Tip stem cells were then compared against the union of myeloid-, MEP-committed cells and LMPPs.

#### Selection of enhancer peak-regions with differentiation-dependent activity

To identify enhancer peak regions with putatively distinct activity between HSC states and cell states of early lineage commitment, we selected enhancer peaks based on their histone marks derived from a published atlas of hematopoietic enhancers^28^. From this atlas we selected enhancers with differentiation state specific H3K4me1 levels. Myeloid enhancers, inactive in stem cells but poised during onset of myeloid differentiation, were defined by low (<25) normalized H3K4me1 CHIP-seq counts in LTs, CLPs and MEPs and high (>50) H3K4me1 levels in CMPs and GMPs. MEP enhancers, were defined by low (<25) H3K4me1 counts in LTs, CLPs, CMPs and GMPs and high (>50) H3K4me1 levels in MEPs. CMPs were not considered as part of the MEP differentiation trajectory because Lara-Astiaso et al.^28,69^ reported that CMPs clustered together with GMPs based on enhancer marks. CLP enhancers, were defined by low (<25) H3K4me1 counts in LTs, CMPs, GMPs and MEPs and high (>50) H3K4me1 levels in CLPs. HSC enhancers, were defined by low (<25) H3K4me1 counts in all progenitor populations but high (>50) H3K4me1 levels in LTs.

#### ChromVar-based quantification of accessibility within sets of marker-loci

To quantify chromatin accessibility in differentiation-associated regions (e.g. promoters/ enhancers with lineage-specific accessibility), we used Signac’s AddChromatinModule^69^ function to compute ChromVar z-scores^47^ for each set of loci. We used ChromVar z-scores because they are—compared to accessibility counts—not confounded by the cell-cycle. This is because ChromVar z-scores are a relative measure of accessibility, quantifying how a cell’s accessibility signal across a given set of loci stands out against background loci in the same cell. Specifically, ChromVar z-scores are computed in two steps. First, for each cell its deviation from the expected accessibility at the loci of interest is computed (expectation based on library size). Then this deviation is converted into a z-score, which quantifies how far it stands out compared to deviations of background loci in the same cell. Background loci are selected to match the loci of interest in terms of GC content and baseline accessibility. Furthermore, we computed ChromVar z-scores based on ATAC matrices containing either only enhancer peaks or only promoter peaks, depending on whether the loci of interests were enhancers or promoters. This ensured that background loci were not only matching the loci of interest in terms of GC content and accessibility but additionally in terms of their type (i.e. promoters/enhancers).

#### Transcriptome-based matching of cells from distinct cell-cycle phases

As illustrated in Figure 5A (left), we selected pairs of adjacent HSPCs (“matched neighbors”) from a 3D RNA-space (computation of PHATE embedding in method section “Construction of RNA-based cell embedding using lineage markers”). Each HSPC-pair contained one cell in G1 phase and another cell in the phase of interest. This one-to-one matching of HSPCs from two distinct cell-cycle phases was done based on a k-nearest neighbor graph which was computed based on the 3D-RNA-embedding using Seurat’s FindNeighbors function. HSPC-pairs were selected from the graph by the following algorithm: Starting from one cell of the first cell-cycle phase, its ten nearest neighbors were screened for the closest cell of the other cell-cycle phase. A match was only accepted if the two cells were each other’s mutual nearest neighbors. We refer to the aggregate of all successfully matched cell pairs as “matched neighbors”. By design, matched neighbors always contain equal amounts of uncommitted hematopoietic progenitors from the two cell-cycle phases. Also, it is enforced that cells from the two cell-cycle phases within the matched neighbors are similarly distributed in the 3D RNA-space used for selection. Matched neighbors were selected for both pairwise comparisons between G1 and the remaining cell-cycle phases (G1-S, G1-G2M). Exemplary, Figure 5B and S8 show that matched G1 and S phase-HSPCs are equally distributed along a differentiation continuum from stem cells to committed progenitors.

